# Necroptotic Signalling Diverts Keratinocyte Fate to Promote Differentiation and Slow Wound Healing

**DOI:** 10.64898/2026.07.13.738083

**Authors:** Holly Anderton, Yingxue He, Natasha Silke, Anisha Lynch-Godrei, Lok Hei Gu, Stacey Brown, Kosuke Shimada, Esther Bandala-Sanchez, Wayne Cawthorne, Shene Chiou, Anne Hempel, Andre L. Samson, James M. Murphy, John Silke

## Abstract

Necroptosis is best known as a lytic, proinflammatory cell-death pathway mediated by RIPK3 and MLKL. Effective wound repair requires the rapid resolution of inflammation, and ongoing necroptotic activity would only exacerbate tissue damage, delaying healing. However, damaged skin presents a trigger-rich environment for necroptotic signalling, an apparent paradox that remains unresolved. Using genetic ablation and pharmacological inhibition across multiple wound models, we show that inhibiting necroptosis accelerates wound closure, revealing that necroptotic signalling normally restrains repair. Surprisingly, we found that MLKL activation in wild-type keratinocytes induces differentiation and membrane repair rather than cell lysis. This adaptive, non-lethal mode of necroptotic signalling preserves barrier integrity but slows re-epithelialisation. Our findings redefine epidermal necroptotic signalling as a stress-responsive program that modulates keratinocyte fate in a trigger-rich environment. Temporarily dampening this pathway may enhance regeneration after barrier loss without compromising immune defence, revealing necroptosis as a tunable mechanism balancing tissue repair and inflammation.

## Introduction

The skin serves as both a physical and immunological barrier, protecting the body by preventing water loss and sensing damage. The skin also mounts rapid immune and repair responses, which are vital for maintaining barrier integrity^1–4^. Following injury, keratinocytes, fibroblasts, and immune cells engage in a highly choreographed inflammatory response to clear debris and restore homeostasis^5,6^. Achieving the right balance is crucial: an inadequate response risks infection, while excessive or prolonged activation can drive tissue damage, leading to repeated cycles of injury and repair that can culminate in chronic, non-healing wounds^6,7^.

Programmed cell death plays a role in this balance between inflammation and repair. Apoptosis promotes resolution through orderly cell clearance^8,9^, whereas necroptosis, a regulated yet lytic process driven by RIPK3 and MLKL, can amplify inflammation and intensify tissue damage^10^.

Necroptosis appears to have first evolved as an anti-pathogen defence strategy, eliminating infected cells when apoptosis was blocked^11^. However, the ability of a wide range of immuno-inflammatory and stress signals to activate the necroptotic pathway suggests that its role has expanded beyond anti-pathogen defence to encompass broader responses to cellular stress^12^.

In reparative settings, a lytic, inflammatory process such as necroptosis seems at odds with tissue recovery and barrier repair. Yet the wound environment is precisely the kind of trigger-rich milieu that favours its activation. Cytokines, damage-associated molecular patterns (DAMPs), and microbial or commensal-derived ligands converge on shared signalling nodes capable of engaging RIPK3 and MLKL^12,13^. This trigger-rich environment provides ample biochemical opportunity for necroptotic activation even in the absence of overt infection, raising the question of how this potentially destructive pathway operates without undermining repair.

We hypothesised that rather than inducing cell death, necroptotic signalling in the wound environment might serve an alternative, adaptive function. To test this, we assessed cutaneous recovery in mice lacking RIPK3 or MLKL and assessed the fate of MLKL-activated keratinocytes using genetic, pharmacological, and live-imaging approaches. We found that necroptotic signalling plays an unexpected, non-lethal role in cutaneous recovery, acting as a stress-adaptive program within the epidermis, rather than a purely cytolytic process. Across multiple models of skin injury, loss of RIPK3 or MLKL accelerates wound closure, demonstrating that necroptotic signalling normally restrains repair. We found that, rather than inducing cell death, MLKL activation in keratinocytes promotes differentiation and membrane repair, diverting cell fate away from lysis, preserving barrier integrity but slowing re-epithelialisation. These findings reveal an unanticipated dimension of necroptosis signalling: a contextually adaptive pathway that integrates inflammatory and metabolic cues to modulate epidermal fate. We propose that necroptosis, long viewed as a destructive endpoint, can also function as a tunable program that balances tissue repair and defence.

## Results

### Necroptotic signalling impedes epidermal recovery in a mouse model of toxic epidermal necrolysis (TEN)

To define the role of regulated cell death during cutaneous injury and repair, we employed the Smac-mimetic (SM)-induced toxic epidermal necrolysis (TEN) model, in which pharmacological IAP antagonism triggers rapid, TNFR1-dependent, keratinocyte death and barrier loss^14,15^. We performed a systematic and comprehensive in vivo genetic dissection using a panel of knockout and mutant mouse strains targeting regulated cell death and inflammatory signalling components to delineate the mechanisms governing SM-induced epidermal injury (Fig. 1A and Table S1).

**Figure 1.**
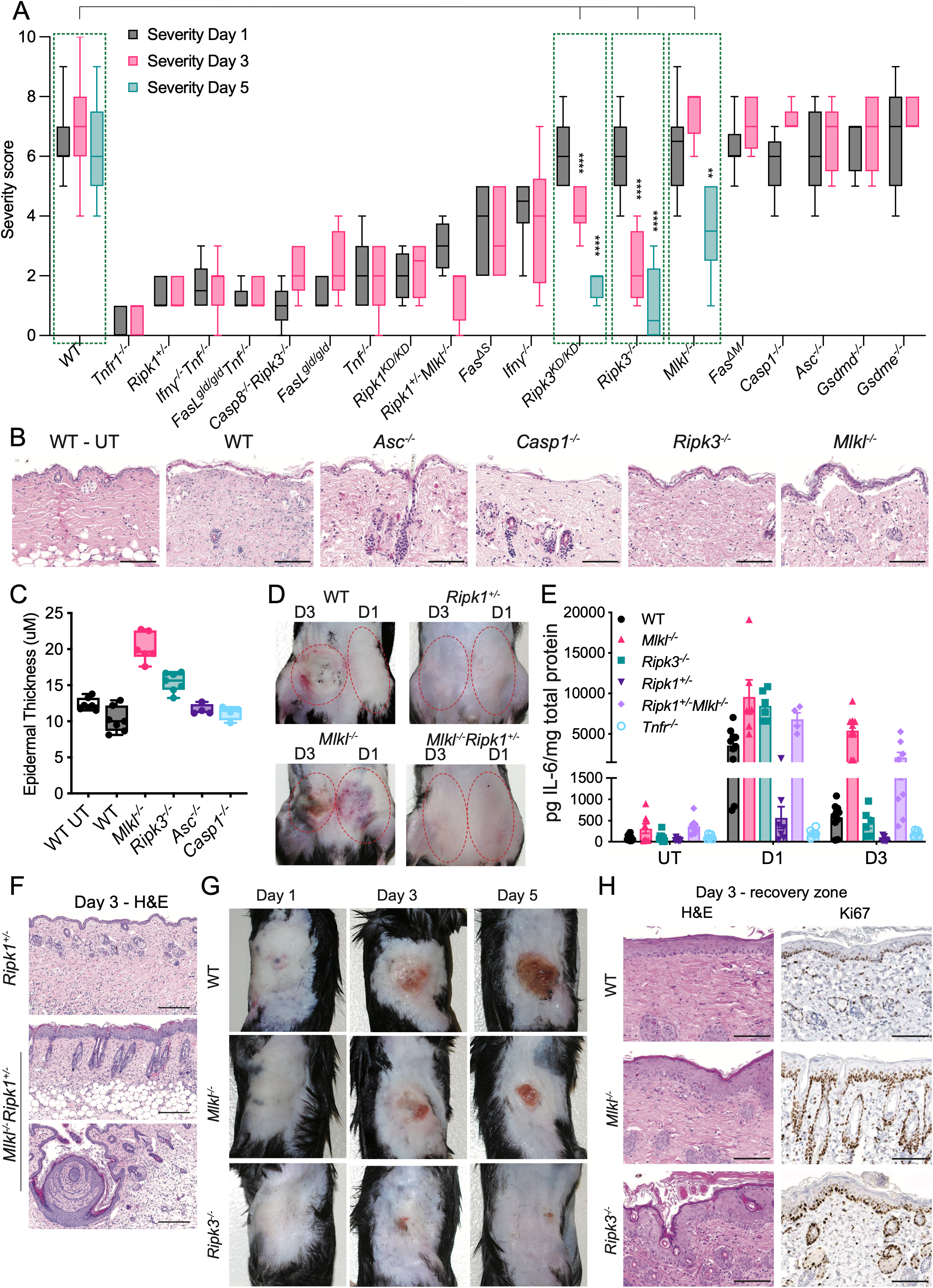
Necroptotic signalling restrains epidermal recovery in a mouse model of toxic epidermal necrolysis (TEN). (A) TEN severity scores following induction by subcutaneous injection of Smac-mimetic (SM) in WT mice and the indicated strains. Severity was scored at days 1 and 3 post-injection for all genotypes, and on day 5 for a subset. n ≥ 4 animals per genotype. KD, kinase-dead allele; Fas^gld/gld^, FasL loss-of-function; Fas^ΔS^, soluble FasL-deficient; Fas^ΔM^, membrane-bound FasL-deficient. Box plots show the interquartile range (25th–75th percentile) with the centre line indicating the median and whiskers representing minimum and maximum values. N ≥ 4 animals per genotype; individual data points are not shown for clarity due to variable sample sizes across genotypes; exact N values and summary statistics are provided in Table S1. (B) Representative H&E-stained skin sections from untreated WT skin (WT-UT) and day-1 TEN lesions from WT and select regulated cell death pathway-deficient mice. Scale = 100μM. (C) Quantification of epidermal remnant thickness at day 1 post-injection for the indicated genotypes. N=4-7, each data point is the average of ≥10 measurements per animal. A box plot shows the median and the interquartile range. (D) Representative images of injection sites in the indicated genotypes at days 1 (right) and 3 (left) following subcutaneous inguinal Smac-mimetic injection. Dashed ovals indicate the lesional sites. (E) IL-6 concentrations in whole skin lysates from the indicated genotypes measured by ELISA assay at untreated baseline (UT), day 1 (D1), and day 3 (D3) following SM injection. Cytokine levels are expressed as pg per mg total protein. N=4-10, each data point is a separate animal. Bars are mean ± SEM. (F) H&E-stained sections of day-3 lesional skin from the indicated genotypes. Scale = 100μM. (G) Representative images of lesions in WT, Mlkl^-/-^, and Ripk3^-/-^mice at days 1, 3, and 5 following SM injection. (H) H&E staining and Ki67 immunohistochemistry of day-3 recovery zones from WT, Mlkl^-/-^, and Ripk3^-/-^ mice. Scale = 100μM.

Consistent with prior work, deletion of Tnfr1 rendered mice largely unresponsive to SM-induced injury, and even Ripk1^+/-^ mice had substantial protection from keratinocyte death and lesion formation^14^. RIPK1 kinase-dead mutants (Ripk1^D138N/D138N^ abbreviated Ripk1^KD/KD^) were similarly protected (Fig. 1A). In contrast, genetic deletion of Tnf or Ifng reduced severity but did not entirely prevent epidermal apoptosis, while mice harbouring a non-functional mutation in FasL (Fas^gld/gld^) exhibited markedly diminished keratinocyte cell death^14^. Extending these findings, we found that combined loss of TNF and IFNγ phenocopied the Fas^gld/gld^ outcome, whereas Fas^gld/gld^Tnf^-/-^ animals were not further protected (Fig. 1A). These findings support a model in which TNF and IFNγ act upstream to prime keratinocytes for FasL-dependent apoptosis^14,16^.

Mice lacking the central inflammasome adaptor ASC, the pyroptotic activator Caspase-1 or the pyroptotic effectors gasdermin D or E, exhibited disease severity and epidermal ablation comparable to wild-type (WT) animals (Fig. 1A and B), indicating that inflammasome activation and pyroptosis do not significantly contribute to SM-induced TEN. Similarly, deletion or mutation of the necroptotic effectors RIPK3 or MLKL did not limit day 1 severity or prevent acute epidermal detachment (Fig. 1A and B), demonstrating that the primary epidermal injury is necroptosis independent. However, despite comparable early injury, Ripk3^−/−^, Ripk3^D143N/D143N^ (kinase dead; abbreviated Ripk3^KD/KD^) and Mlkl^−/−^ mice showed accelerated recovery at later timepoints, with reduced severity evident by day 5 (Fig. 1A). This suggested that necroptotic signalling does not initiate epidermal loss in this model, but appears to restrain epidermal repair upon barrier disruption.

Notably, this altered epidermal response was already apparent at day 1, before overt differences in lesion resolution. The remnants of separated epidermis visible on day 1 were modestly thicker in Ripk3^−/−^ and Mlkl^−/−^ mice than in WT and other non-rescuing genotypes (Fig. 1B, C and S1A). Although keratinocyte loss was substantial across all strains, this histological distinction suggested that the absence of necroptotic effectors may subtly alter epidermal dynamics in barrier-disrupted skin, even at an early stage.

To further explore early events following SM injection, we prepared an IHC time course on WT samples. We observed early epidermal proliferation (Ki67) starting one hour post-injection, conspicuously before the onset of apoptosis (Fig. S1B). The earliest T-cell and general immune cell infiltration (CD3 and CD45, respectively) occurred concurrently with the increase in Ki67 at 1 hour; however, a notable escalation in these markers correlated with elevated cleaved caspase-3 (CC3) levels from 3 to 6 hours post-injection (Fig. S1B).

Ripk1 heterozygosity prevented lesion formation but induced mild oedema at the injection site, with a moderate increase in IL-6 at day 1 compared to untreated controls, and Ripk1^KD/KD^ mice were similarly protected (Fig. 1A, D and E). Lesion formation remained absent in Ripk1^+/-^Mlkl^-/-^ mice (Fig. 1D), yet, intriguingly, IL-6 was markedly more elevated than in Ripk1^+/-^ or WT mice at the same time point (Fig. 1E). IL-6 levels remained high on day 3 in both the Mlkl^-/-^ and the Ripk1^+/-^Mlkl^-/-^ mice even as inflammation resolved in WT and Ripk1^+/-^ mice (Fig. 1E). In contrast, CCL2 levels were lower in Mlkl^-/-^ and the Ripk1^+/-^Mlkl^-/-^ mice than WT and Ripk1^+/-^ mice (Fig. S1C). Additionally, Ripk1^+/-^Mlkl^-/-^ mice exhibited moderate epidermal hyperplasia and developed keratin pearl structures by day 3 post-injection, indicative of significant and rapid epidermal hyperproliferation (Fig. 1F).

Despite similar severity and extensive epidermal ablation in Ripk3^-/-^ mice at day 1, their TEN lesions were significantly improved by day 3 compared with WT mice. This was also the case for RIPK3 kinase-dead mice, which are unable to phosphorylate and activate MLKL^17^. Although the accelerated recovery was not initially evident in Mlkl^-/-^ mice on day 3, by days 5 and 7 they also showed a marked reduction in lesion size and severity compared to WT mice (Fig. 1A, G and S1D). H&E and Ki67 staining on day 3 post-injection revealed a thicker epidermis and higher proliferation rates in the healing zones of both MLKL-and RIPK3-deficient mice (Fig. 1H), suggesting a rapid cellular response to epidermal injury that may account for their accelerated recovery dynamics.

Bioplex analysis of additional cytokines and chemokines at day 1 post-injection further illustrates the unique inflammatory profiles of Mlkl^-/-^ and Ripk3^-/-^ mice in this model. Cytokines such as MIP1b/CCL4, IL-5, and IL-10 were elevated in both Mlkl^-/-^ and Ripk3^-/-^ mice compared to WT controls, while others, such as G-CSF and IL-17, were specifically increased in Mlkl^-/-^ mice (Fig. S1E). Together, these data indicate that deficiency of RIPK3 or MLKL does not suppress the inflammatory response but instead reshapes it.

### Necroptotic Knockout Mice Have Reduced Dermal Injury Following SM Injection

Although the primary SM-induced injury is necroptosis-independent epidermal apoptosis, there is also dermal damage, which appears reduced in mice lacking necroptotic effectors. By day 7 post-injection, histological examination showed that the proliferative healing edges in the epidermis of WT mice extended beneath an area of dermal necrosis and cellular debris (eschar; indicated by a green dotted line, Fig. 2A). In contrast, at this time, Ripk3^-/-^ mice exhibited nearly complete epidermal coverage, with only residual signs of epidermal hyperplasia. Mlkl^-/-^ mice had more advanced recovery than WT controls, but less than Ripk3^-/-^ mice, with re-epithelialisation still ongoing under patches of dead epidermis. Importantly, the healing edges in these knockout mice did not extend as deeply into the dermis, reflecting significantly less dermal damage.

**Figure 2.**
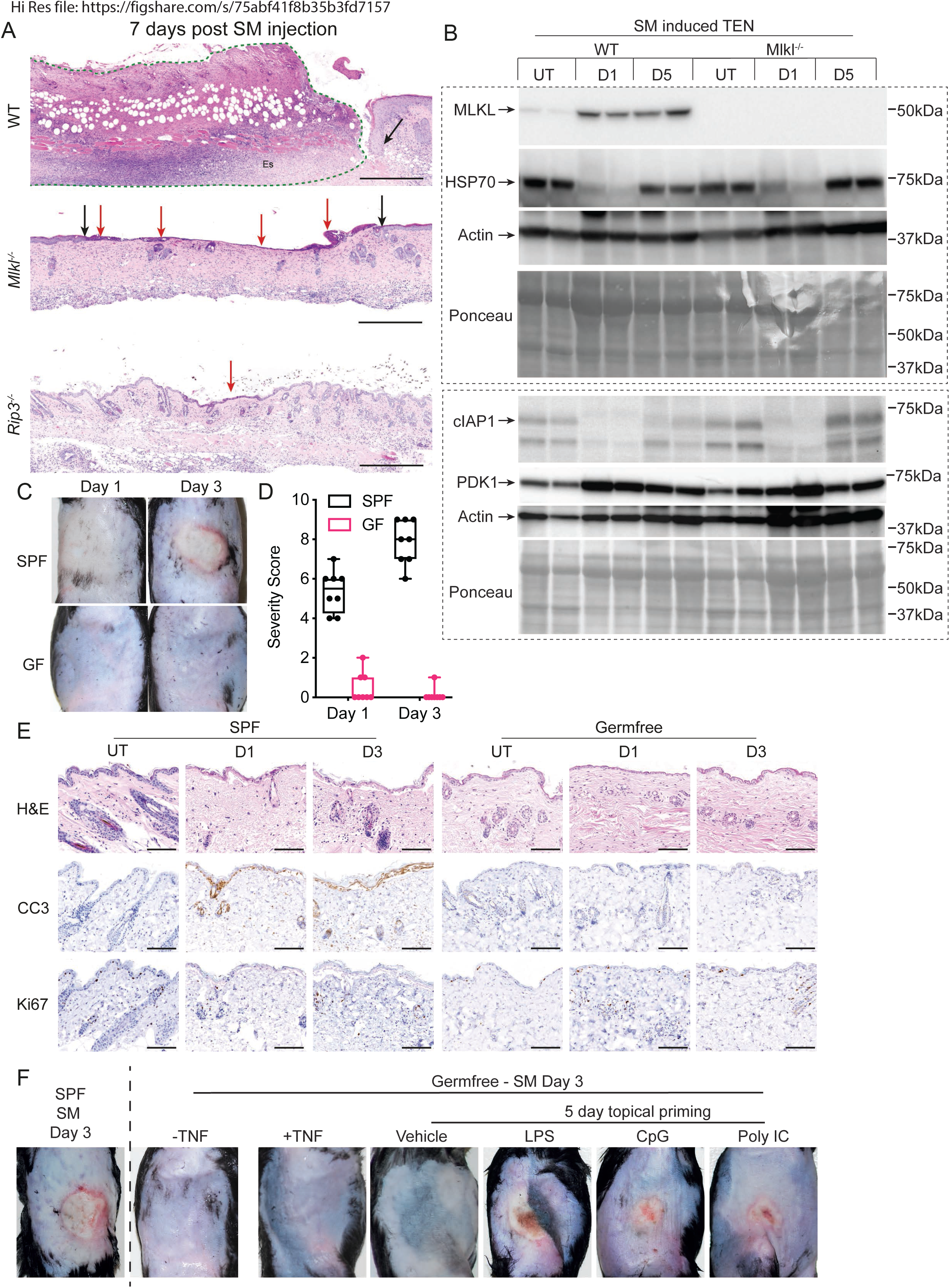
Reduced dermal injury and altered signalling responses in necroptotic knockout mice following SM injection. (A) H&E-stained skin sections 7 days after Smac-mimetic (SM) injection in WT, Mlkl^-/-^, and Ripk3^-/-^ mice. The dashed green line outlines the region of dermal eschar (Es) and cellular debris underlying the regenerating epidermis in WT skin. Arrows indicate epidermal healing fronts. Scale = 500μM. (B) Immunoblot analysis of whole skin lysates from WT and Mlkl^-/-^ mice at untreated baseline (UT), day 1 (D1), and day 5 (D5) following SM injection. Blots as indicated. Ponceau and actin staining serve as a loading control. (C) Representative images of injection sites in specific pathogen–free (SPF) and germ-free (GF) mice at days 1 and 3 following SM injection. Scale = 5mm. (D) TEN severity scores in SPF and GF mice at days 1 and 3 following SM injection. Box plots show the interquartile range (25th–75th percentile) with the centre line indicating the median and whiskers representing minimum and maximum values. Each point represents an individual animal. (E) H&E staining and immunohistochemistry for cleaved caspase-3 (CC3) and Ki67 in skin from SPF and GF mice at UT, D1, and D3 following SM injection. Scale = 100μM (F) Representative images from germ-free mice on day 3 following SM injection with or without TNF co-injection, or following 5 days of topical priming with vehicle, LPS, CpG, or Poly(I:C) prior to SM challenge.

One possible explanation for the increased dermal injury in WT mice is that necroptosis is a secondary effect triggered by SM-induced IAP inhibition and loss in the dermis. Western blot analysis of whole skin lysates over a time course revealed an increase in total MLKL in WT mice at day 1, which persisted until at least day 5, after IAP levels had rebounded (Fig. 2B).

Additionally, a notable increase in PDK1 was observed in both WT and Mlkl^-/-^ samples by day 1 (Fig. 2B). This protein plays a critical role in activating glycolysis, contributing to pro-proliferative and pro-migratory roles in wound healing^18^; however, it has also been shown to have caspase-8 inhibition activity^19^, potentially providing an in vivo mechanism that enables necroptotic signalling in this context.

To assess whether microbially-induced necroptosis contributes to the slower recovery observed in WT mice relative to mice deficient in necroptotic effectors, we tested the SM-induced TEN model in axenic or germ-free (GF) mice. Remarkably, 8 of 10 GF mice showed no measurable reaction to SM injection (Fig. 2C and D), and, critically, cleaved caspase-3 staining revealed an absence of keratinocyte apoptosis and epidermal disruption, while Ki67 staining indicated no increase in epidermal proliferation (Fig. 2E). The few GF mice that exhibited any response showed only mild localised swelling that did not progress to lesion formation (Fig. S2A). Cleaved caspase-3 staining from day 1 post-SM in these mildly responsive mice revealed the minimal nature of these reactions (Fig. S2A). Therefore, a GF environment, rather than reducing an inflammatory reaction by preventing microbial access to the damaged skin, prevented the response and lesion formation entirely. Co-injection of SM with TNF was insufficient to sensitise GF mice to SM-induced lesions; however, topical application of the microbial ligands LPS, PolyIC, or CpG prior to SM injection elicited a lesional response (Fig. 2F). These findings indicate that microbial products generate a SM-responsive environment in the skin.

### MLKL delays epidermal recovery in a keratinocyte-specific model of barrier disruption

The absence of a lesional response in GF mice precluded the use of the SM model to assess whether microbially induced necroptosis contributed to the healing advantage observed in Mlkl^-/-^ and Ripk3^-/-^ mice. Additionally, we remained unsure whether the hyperinflammation and hyperproliferation observed in necroptotic knockout mice were a direct consequence of the SM itself. Therefore, to investigate accelerated healing in Mlkl^-/-^ mice, we explored alternative models of cutaneous disruption, starting with tamoxifen-inducible cFLIP epidermal knockout mice (Cflar^fl/fl^Tg(K14-creERT2), hereafter cFlip^k14ERcre^)^20,21^. Topical application of tamoxifen leads to keratinocyte-specific loss of the caspase-8 inhibitor cFLIP. Depletion of cFLIP, therefore, leads to keratinocyte apoptosis and loss of the epidermal layer. Unlike the SM injection model, other skin-localised cells are unaffected. We crossed cFlip^k14ERcre^ mice to MLKL-deficient mice to examine recovery following induced epidermal cFLIP deletion and subsequent barrier destruction.

As expected, MLKL deficiency had no impact on the apoptotic injury triggered by cFLIP loss, with the entire tamoxifen-treated area visibly and equally impacted in WT and Mlkl^-/-^ mice (Fig. 3A). However, by day 14, ten days after the last tamoxifen application, epidermal recovery in Mlkl^-/-^ mice was significantly more advanced than in WT counterparts, illustrating the dramatic differences in healing dynamics (Fig. 3B). Consistently, histological analysis of the time course revealed that both WT and Mlkl^-/-^ mice displayed similar characteristics at the early stages. At day 4, increased epidermal proliferation (Ki67) and apoptosis (CC3) were noted equally in both strains (Fig. 3C). Immune cell recruitment, including numerous neutrophilic micro-abscesses along the disrupted epidermis (MPO, Fig. S3A) and a robust macrophage presence throughout the dermis (F480, Fig. S3A), was comparable in WT and Mlkl^-/-^ mouse skin across the time course. However, from day 7 and, more notably, by day 9, differences in healing became apparent, with Mlkl^-/-^ mice showing more extensive re-epithelialisation under the scab layers. By day 11, the recovered epithelium of Mlkl^-/-^ mice was notably thicker than equivalently recovered areas in the WT mice, correlating with increased basal keratinocyte proliferation in the knockouts (Ki67, Fig. 3C). In contrast to the SM-induced TEN model, Mlkl^-/-^ mice trended towards a mildly reduced inflammatory cytokine profile across the time course compared to WT mice (Fig. S3B). As with the first model, there was an overall increase in PDK1 in both WT and Mlkl^-/-^ mice upon barrier damage (Fig. S3C).

**Figure 3.**
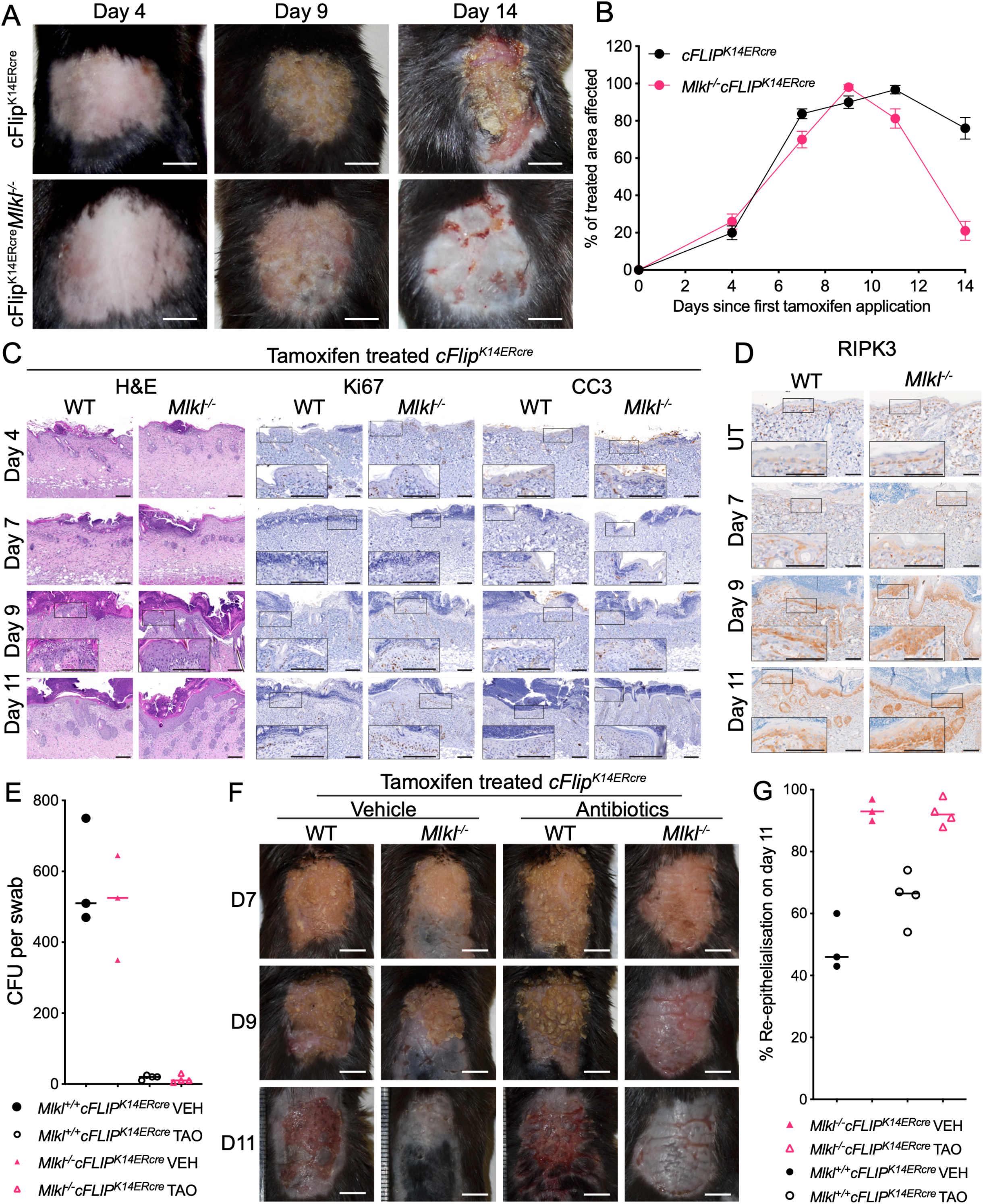
Epidermal recovery is delayed by MLKL in a keratinocyte-specific model of barrier disruption. (A) Representative images of cFLIP-depleted skin in cFlip^K14ERcre^ and Mlkl^-/-^cFlip^K14ERcre^mice at days 4, 9, and 14 following the first tamoxifen application. Scale = 6mm (B) Quantification of the percentage of the treated area affected over time following tamoxifen-induced cFLIP deletion in cFlip^K14ERcre^ and Mlkl^-/-^cFlip^K14ERcre^ mice. N=10, data points are mean ± SEM. (C) Histological analysis of skin during the recovery time course. Representative H&E staining and immunohistochemistry for Ki67 and cleaved caspase-3 (CC3) at days 4, 7, 9, and 11 following the start of tamoxifen treatment. Scales = 100μM. (D) Immunohistochemistry for RIPK3 in untreated skin and during recovery (days 7, 9, and 11 from start of tamoxifen) from mice of the indicated genotypes. Scales = 100μM. (E) Quantification of bacterial colony-forming units (CFU) recovered from skin swabs collected at day 7 post tamoxifen in vehicle (VEH) or triple antibiotic ointment (TAO) treated animals of the indicated genotypes. N=3-4. (F) Representative clinical images of tamoxifen-treated skin from mice treated with vehicle or TAO throughout. Images are shown at days 7, 9, and 11 following the first tamoxifen application. Scabs were removed for the day 11 pictures so that the extent of epidermal recovery is not obscured. Scale = 6mm. (G) Quantification of the percentage of re-epithelialisation at day 11 in VEH or TAO-treated animals of the indicated genotypes. N=3-4, each point represents an individual animal.

Interestingly, immunostaining for total RIPK3 showed expression in untreated epidermis, specifically in basal keratinocytes, in both WT and Mlkl^-/-^ mice, with expression lost in suprabasal layers. This pattern is also observed in the recovering epidermis of both groups, suggesting that RIPK3 expression marks basal keratinocytes, indicating that the capacity to undergo necroptosis is likely reduced as keratinocytes differentiate (Fig. 3D).

To ascertain whether microbially driven cutaneous necroptosis was responsible for the effect, we treated cFLIP^K14ERcre^ mice with topical antibiotics concurrently and following tamoxifen-induced cFLIP deletion. Swabbing the treatment area on day 7, we found a substantial reduction in colony forming units (CFU) present on the skin of the antibiotic-treated animals of both genotypes, confirming the general effectiveness in reducing the commensal load in the skin (Fig. 3E). While the use of topical antibiotics and the reduction in microbial numbers appears to have resulted in a mild improvement in re-epithelialisation of WT mice by day 11, the treatment did not phenocopy the accelerated healing of Mlkl^-/-^ mice (Fig. 3F and G).

### The cutaneous healing advantage also occurs in mechanical wounds

Our results suggested that loss of the epidermal layer activates the necroptotic pathway and delays healing, an effect that, surprisingly, is independent of concurrent microbial exposure. We then tested whether the healing advantage observed in necroptosis-deficient mice was also evident in mechanical wounds using excision wound studies. And indeed, wounds in Ripk3^-/-^ and Mlkl^-/-^ mice exhibited faster closure than in WT mice, with day 6 wounds significantly smaller (Fig. 4A and B). RIPK3 kinase-dead mice also showed accelerated wound closure. Similarly, RIPK1 kinase-dead mutants, which are resistant to TNF-induced necroptosis^22^, exhibited advanced wound closure comparable to other knockout strains (Fig. 4A and B).

**Figure 4.**
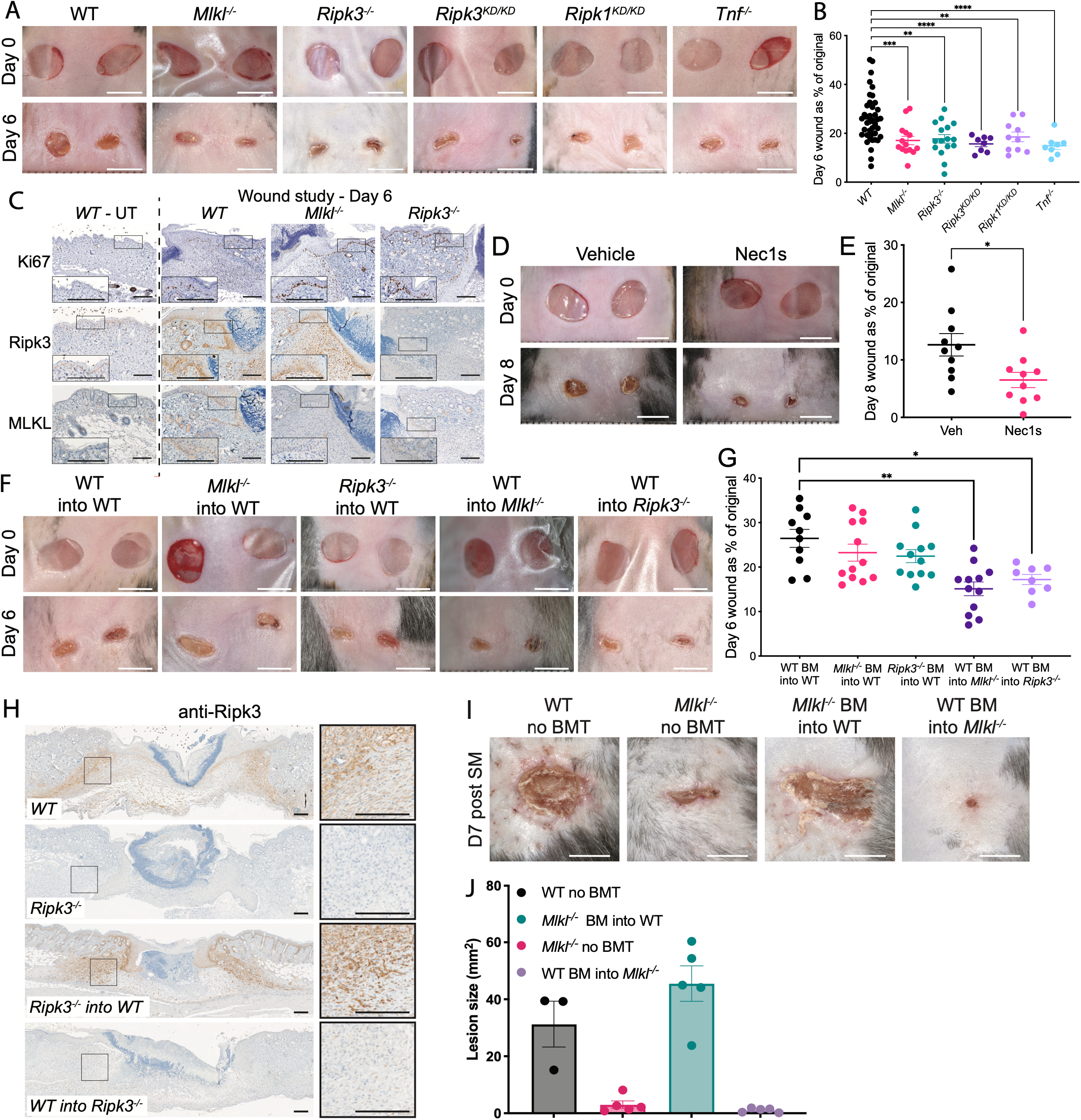
Accelerated wound healing in necroptosis-deficient mice and pharmacologic inhibition of necroptosis. (A) Representative images of excisional wounds in the indicated strains at day 0 and day 6 following wounding. Scale = 5mm. KD = Kinase dead; Ripk3^KD/KD^ are Ripk3^D143N/D143N^, Ripk1^KD/KD^are Ripk1^D138N/D138N^. (B) Quantification of wound size at day 6 expressed as a percentage of the original wound area for the indicated genotypes. (C) Immunohistochemical analysis of wound edges at day 6 showing Ki67, RIPK3, and MLKL staining in the indicated genotypes. Untreated WT skin (WT-UT) is shown for comparison. Scale = 200μM. (D) Representative images of excisional wounds in WT mice treated with vehicle or the RIPK1 inhibitor Nec-1s at day 0 and day 8 following wounding. Scale = 5mm. (E) Quantification of wound size at day 8 expressed as a percentage of the original wound area in vehicle-and Nec-1s-treated animals. (F) Representative images of excisional wounds in bone marrow transplantation (BMT) chimeras at day 0 and day 6 following wounding. Scale = 5mm. (G) Quantification of wound size at day 6 expressed as a percentage of the original wound area in the indicated BMT chimeras. (H) Immunohistochemistry for RIPK3 in day-6 wounds from the indicated BMT groups. (I) Representative images of SM-induced TEN lesions at day 7 in the indicated BMT groups. Scale = 5mm. (J) Quantification of SM induced lesion size at day 7 in the indicated BMT groups. Graphs show mean ± SEM. Each point represents an individual wound. Mice typically received two wounds, but wounds in which the dressing was disrupted, resulting in premature exposure, were excluded from quantification. Significance was calculated using an unpaired t-test with Welch’s correction. P < 0.05 (*), P < 0.01 (**), P < 0.001 (***), P < 0.0001 (****).

Histology of the wound edges of WT, Ripk3^-/-^ and Mlkl^-/-^ mice showed increased proliferation (Ki67) in the knockout mice near the wound edge (Fig. 4C). Notably, there was again clear, specific expression of RIPK3 in both WT and Mlkl^-/-^ basal keratinocytes (Fig. 4C). This was more pronounced in day 6 wounds compared to untreated WT skin, but there was no obvious difference in RIPK3 expression between WT and Mlkl^-/-^ at day 6. There was also strong RIPK3 expression in the regenerating dermis of both WT and Mlkl^-/-^ mice, especially at the interface between the granulation tissue and the remodelling dermis.

MLKL expression was less apparent generally and was undetectable in untreated WT skin; however, specific MLKL expression in basal WT keratinocytes can be seen in regenerating epithelia near the healing site (Fig. 4C). An increase in dermal expression of total MLKL is not immediately obvious. Still, closer examination of the dermal regeneration area, where RIPK3 was upregulated, reveals specific MLKL staining as well (Fig. S4A). These results are consistent with Western blot data, which showed low total MLKL in untreated skin but increased expression following wounding (Fig. S4B). Once again, we observed increased PDK1 in WT and Mlkl^-/-^ mouse wounds, particularly at day 6 relative to day 0 and a generally higher level in Ripk3^-/-^ mice (Fig. S4C). Our findings from genetic disruption of necroptosis effectors were phenocopied by pharmacological inhibition of the pathway in WT mice with the RIPK1 inhibitor Necrostatin-1s (Nec-1s). Wounds treated with Nec-1s showed markedly improved healing compared to those treated with vehicle alone (Fig. 4D and 4E).

### The healing advantage conferred by necroptosis pathway deficiency is a tissue-localised effect

We next conducted bone marrow transplantation experiments to discern if the healing advantage was mediated by tissue-specific or immune cell-related mechanisms. Mlkl^-/-^ and Ripk3^-/-^ bone marrow was transplanted into WT mice, and vice versa, prior to wound induction. We found that WT mice with Mlkl^-/-^ or Ripk3^-/-^ bone marrow did not exhibit enhanced healing compared to controls, suggesting that the healing advantage is intrinsic to the skin-localised cells rather than the haematopoietic compartment (Fig. 4F and 4G). Conversely, Mlkl^-/-^ and Ripk3^-/-^ mice transplanted with WT bone marrow maintained their healing advantage, underscoring a tissue-specific effect (Fig. 4F and 4G). These experiments also clarified that the majority of the strong dermal RIPK3 expression in WT and Mlkl^-/-^ wounds is tissue-localised because WT mice receiving Ripk3^-/-^ bone marrow had a similar pattern of RIPK3 dermal expression as untransplanted WT mice (Fig. 4H). There are scattered RIPK3-positive cells in the wound area of Ripk3^-/-^ mice receiving WT bone marrow, confirming RIPK3 expression in infiltrating bone marrow-derived cells, but it is clear that the primary effect is tissue-specific.

We obtained similar results with bone marrow transplantation prior to SM-induced TEN. WT mice that received Mlkl^-/-^ bone marrow showed no improvement in recovery speed compared to untransplanted WT mice, whereas Mlkl^-/-^ mice that received WT bone marrow not only retained their healing advantage but also exhibited more complete resolution at day 7 than untransplanted Mlkl^-/-^mice (Fig. 4I and J).

### Cultured keratinocytes are resistant to necroptotic cell death

Although our MLKL and RIPK3 staining show that both epidermal and dermal healing fronts express the necroptotic machinery, our comparative analyses across all three models suggests the importance of the epidermis in the healing response. In particular, faster healing, necroptosis-deficient mice consistently displayed heightened keratinocyte proliferation and a markedly thicker repairing epidermis. Meanwhile, there was no evidence of excessive cell death or necrotic keratinocytes in WT mice. This suggests that altered epidermal dynamics, rather than loss of keratinocytes, may be a major contributor to the slower cutaneous repair observed in WT mice compared with those lacking necroptotic effectors.

To explore whether keratinocytes can divert necroptotic signalling toward a non-lytic fate, we turned to immortalised human keratinocytes (N/TERT). The in vivo wound environment contains multiple potential, but undefined, upstream inputs that may contribute to RIPK3– MLKL engagement. Our in vitro system was therefore not intended to reproduce the physiological activation of the necroptosis pathway. Instead, treatment with TNF, Smac-mimetic, and IDN-6556 (TSI) was used to activate the pathway in vitro, allowing us to examine how keratinocytes respond once MLKL is engaged. We nevertheless treated cultured keratinocytes with a range of TSI concentrations, hypothesising that the strength of the necroptotic stimulus experienced by keratinocytes in vivo could be weaker than that typically employed in vitro. Therefore, in addition to the standard condition used for cell death assays (TSI-H; TNF 100 ng/mL, SM 250 nM, IDN 5 μM), we included low (TSI-L; TNF 4 ng/mL, SM 20 nM, IDN 2.5 μM) and medium (TSI-M; TNF 25 ng/mL, SM 50 nM, IDN 2.5 μM) concentration conditions.

Compared to HT29s, which were very sensitive to necroptotic death even at the lowest concentration of TSI tested, N/TERT keratinocytes did not undergo substantial cell death, with a small increase in the proportion of PI-positive cells seen only with the highest TSI dose (Fig. 5A and B). While N/TERT cells were not sensitive to TSI-induced death, Western blotting of 3 independent wild-type and 5 independent MLKL^-/-^ keratinocyte lines confirmed specific MLKL activation loop phosphorylation at S357/T358, a mark of pathway activation (Fig. 5C). On the other hand, cell growth was notably arrested in TSI-treated WT N/TERT cells, in a dose-dependent manner, compared to untreated cells (Fig. 5D) and single-stimulus controls (Fig. S5A). WT TSI-treated N/TERT cells also exhibited distinct morphological changes, becoming larger and flatter, resembling differentiating keratinocytes in culture (Fig. 5E).

**Figure 5.**
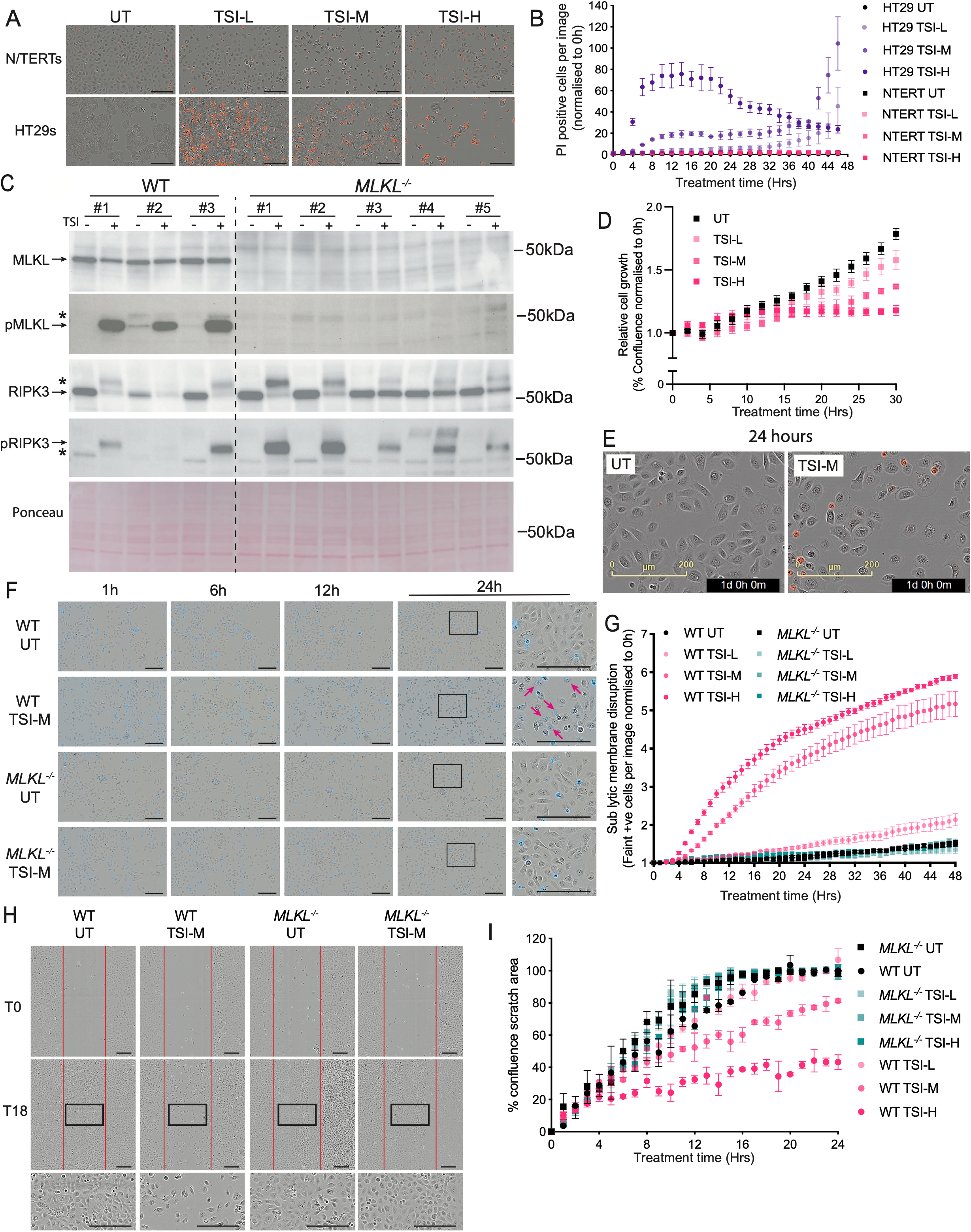
Cultured keratinocytes are resistant to necroptotic cell death. (A) Representative images of PI uptake in N/TERT keratinocytes and necroptosis-sensitive HT29 cells following treatment with TNF + SMAC mimetic + IDN (TSI) at low (TSI-L), medium (TSI-M), or high (TSI-H) concentrations. Untreated (UT) cells are shown for comparison. Scale = 200μM. (B) Quantification of PI-positive cells over time following treatment with TSI-L (low dose), TSI-M (medium dose), or TSI-H (high dose) in N/TERT and HT29 cells. n = 2 technical replicates except NTERT TSI-H (n=4) and HT29 TSI-H (n=3). (C) Immunoblot analysis of independent clones of WT and MLKL^-/-^ N/TERT cells following TSI treatment. Ponceau staining serves as a loading control. (D) Relative cell growth measured over time in WT N/TERT cells following treatment with TSI-L, TSI-M, or TSI-H. N = 3 independent experiments (n=3 wells per experiment). (E) Representative phase-contrast images of WT N/TERT cells untreated (UT) or treated with TSI-M (24-hour timepoint). Scale= 200μM. (F) Time-course images of WT and MLKL^-/-^ N/TERT cells following UT or TSI-M treatment showing faint uptake of cell-permeability dye at 0, 6, 12, and 24 hours post-treatment. Scale = 200μM (G) Quantification of faint dye-positive cells over time in WT and MLKL^-/-^ N/TERT cells treated with UT, TSI-L, TSI-M, or TSI-H (n = 5 technical replicates). (H) Representative images of scratch assays in WT and Mlkl^-/-^ N/TERT cells treated with UT or TSI-M at time 0 (T0) and 18 hours (T18). Scale = 200μM. (I) Quantification of scratch wound closure measured as percentage confluence of the scratch area over time in WT and Mlkl^-/-^ N/TERT cells, UT, or treated with TSI-L, TSI-M, or TSI-H (n=3 technical replicates except Mlkl^-/-^ UT, which had n=2). Data on graphs are shown as mean ± SEM.

While examining these morphological changes over time, we also noted an increase in WT N/TERT cells that were faintly positive for cell permeability dyes upon TSI treatment. These cells were not dead and exhibited a differentiated morphology at later timepoints. This mild dye uptake was dose-dependent in TSI-treated WT N/TERT cells and was not evident in either untreated WTs or in TSI-treated MLKL^-/-^ N/TERT cells (Fig. 5F and G).

This suggests that necroptotic signalling, rather than inducing cell death, promotes differentiation and consequently reduces keratinocyte proliferation. To test this, we used a scratch wound assay and, consistent with our in vivo observations, found that TSI-treated WT N/TERT cells had a dose-dependent delay in healing compared to untreated WT and to both untreated and TSI-treated MLKL^-/-^ N/TERT keratinocytes (Fig. 5H and I).

This differentiation effect was not specific to TNF-induced necroptosis. N/TERT cells were similarly resistant to ZBP1-mediated necroptosis induced by CBL0137^23^ in combination with IFNγ priming and caspase inhibition (Fig. S5B and C). As with the TSI treatment, this necroptotic stimulus was sufficient to cause substantial MLKL-dependent cell death in HT29s, but not N/TERT cells (Fig. S5B), while inducing morphological changes in WT N/TERT keratinocytes consistent with a differentiating profile (Fig. S5C).

### MLKL promotes early keratinocyte differentiation at sites of epidermal regeneration

In healthy skin, an increasing calcium gradient in the epidermis promotes keratinocyte differentiation and upward migration^24^, and this gradient is lost in damaged skin^25^. Consistent with keratinocyte sensitivity to increasing Ca^2+^ levels, increasing extracellular calcium is sufficient to induce expression of differentiation markers in vitro^24,26^. Given that activated MLKL is known to affect membrane permeability and ion flu^27,28^, and our observation that TSI-treated WT keratinocytes show signs of sublethal plasma membrane disruption, we hypothesised that MLKL activation can trigger calcium flux, thereby initiating the differentiation program in keratinocytes.

Accordingly, we examined early (keratin 1, KRT1) and late (filaggrin, FLG) differentiation markers in WT and Mlkl^-/-^ skin across the three models of epidermal injury and regeneration. In the SM-induced TEN model, WT epidermis exhibited a pronounced zone of KRT1 expression preceding the onset of contiguous FLG expression in the upper epidermis, extending from the proliferative zone towards the healing edge (Fig. 6A). In contrast, Mlkl^-/-^ epidermis showed a marked reduction in this early differentiation zone, with minimal KRT1 detected between the proliferative front and regions of established FLG expression. Quantification of the distance between the onset of KRT1 and FLG expression confirmed a contraction of this intermediate differentiation zone in Mlkl^-/-^ mice (Fig. 6B).

**Figure 6.**
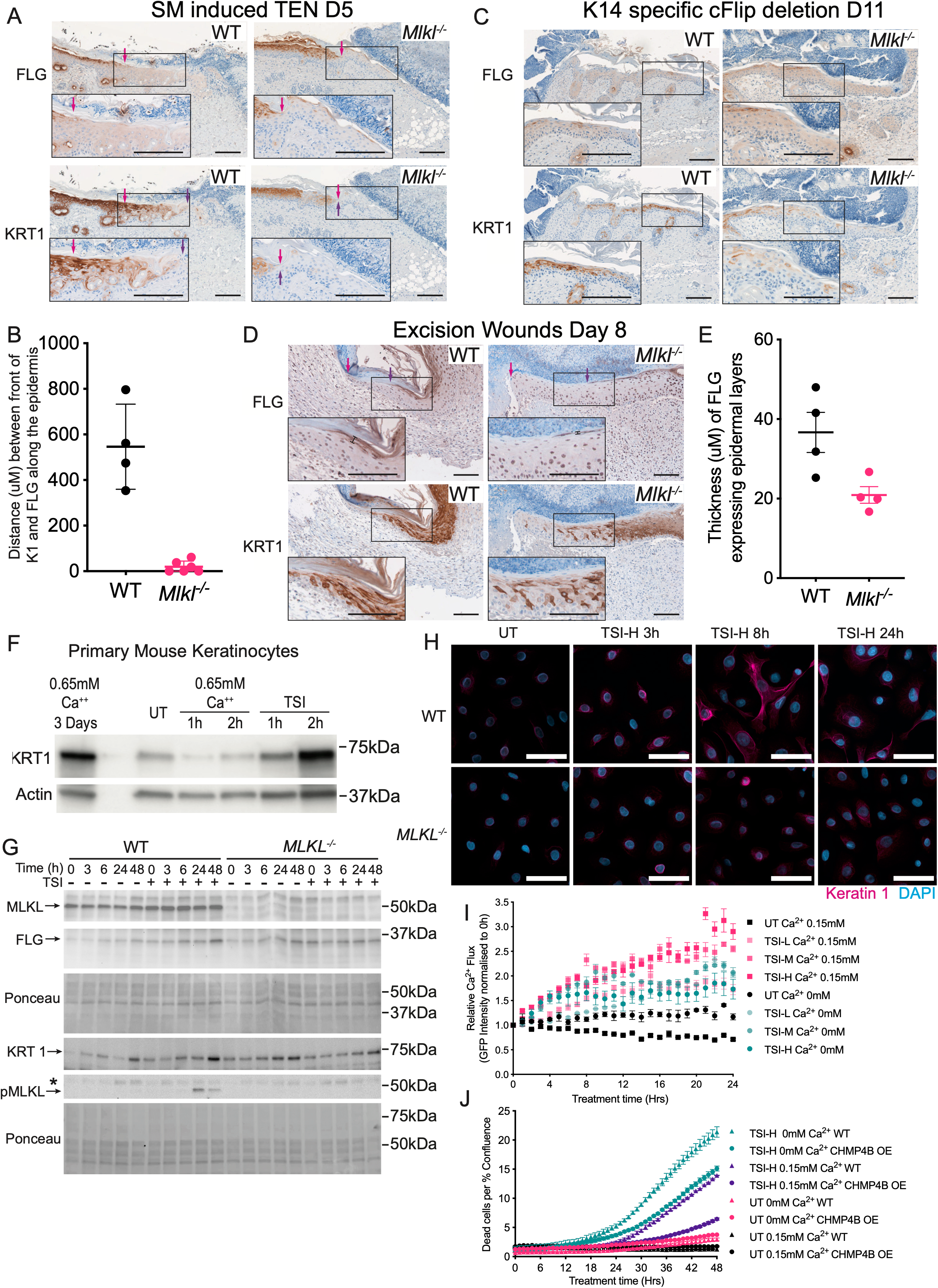
MLKL-dependent signalling promotes keratinocyte differentiation and calcium flux. (A) Representative immunohistochemistry of Keratin 1 (KRT1) and Filaggrin (FLG) in WT and Mlkl^-/-^ epidermis at day 5 following SM-induced TEN. Adjacent sections were used for KRT1 and FLG staining. Arrows indicate the onset of contiguous KRT1 (purple) and FLG (magenta) expression used for spatial comparison. Scale = 100μM. (B) Quantification of the distance (µm) between the onset of KRT1 and FLG expression along the epidermis in WT and Mlkl^-/-^ skin from the SM model. Each data point is a separate mouse, N=4 (WT) or 6 (Mlkl^-/-^). (C) Representative K1 and FLG staining in WT and Mlkl^-/-^ epidermis at day 11 following K14-specific cFLIP deletion. Regions shown correspond to areas with comparable emergence of FLG expression. Scale = 100μM. (D) Representative K1 and FLG staining at the wound edge in WT and Mlkl^-/-^ mice at day 8 following excisional wounding. Arrows indicate the healing front (purple) and FLG (magenta). Insets show higher magnification of boxed regions. Black bars indicate the location of FLG thickness measurements. Scale = 100μM. (E) Quantification of the thickness (µm) of FLG-positive epidermal layers measured 100 µm from the onset of contiguous FLG expression in WT and Mlkl^-/-^ wounds. Each data point is a separate mouse, N=4. (F) Immunoblot analysis of Keratin 1 expression in primary mouse keratinocytes treated with a standard cell death assay dose of TNF + SMAC mimetic + IDN (TSI) or high-calcium medium. Cells were collected at the indicated time points following stimulation. (G) Immunoblot analysis of Keratin 1 and filaggrin expression in WT and MLKL^-/-^ N/TERT keratinocytes following treatment with TSI-M over the indicated time course. (H) Representative confocal images of WT and MLKL^-/-^ N/TERT keratinocytes stained for Keratin 1 (magenta) and DAPI (blue) following treatment with TSI. Scale = 50μM. (I) Calcium flux in gCaMP8-expressing WT N/TERT keratinocytes following treatment with increasing concentrations of TSI in standard medium (0.15 mM Ca²⁺) or calcium-free medium. Fluorescence intensity was quantified over time as a measure of cytoplasmic calcium. Representative experiment shown with n=3 technical replicates. (J) Quantification of cell death in WT N/TERT cells with or without CHMP4B overexpression (OE) following treatment with high-dose TSI in standard (0.15mM Ca²⁺) or calcium-free medium. Representative experiment shown with n=3 technical replicates. Data in graphs are shown as mean ± SEM.

In the K14-specific cFLIP deletion model, epidermal regeneration is less spatially ordered, precluding robust quantification. However, when comparing regions with equivalent emergence of FLG expression, WT epidermis consistently exhibited a higher density of KRT1-positive cells than Mlkl^-/-^ tissue (Fig. 6C). This supports a similar role for MLKL in promoting early differentiation, despite differences in tissue architecture and regenerative dynamics.

In excisional wounds, the effect of MLKL deficiency on KRT1 expression patterns was again evident with Mlkl^-/-^ epidermis displaying a reduced density of KRT1-positive cells in the region between the migrating front and areas of FLG expression (Fig. 6D). In addition, compared to WT, Mlkl^-/-^ epidermis exhibited a reduced vertical thickness of FLG-positive layers when measured a short distance (100 µm) back from the onset of contiguous FLG expression, consistent with diminished maturation of the upper epidermis (Fig. 6E). The distance between the wound edge and the onset of FLG expression also appeared extended in Mlkl^-/-^ mice, although technical variability in tissue orientation and sectioning depth limited formal quantification.

Taken together, these findings support a model in which MLKL promotes early keratinocyte differentiation during epidermal regeneration. While Mlkl^-/-^ mice are ultimately capable of restoring a stratified epidermis, the spatial dynamics of differentiation are altered, with a reduced and delayed early differentiation program near sites of barrier disruption. This is consistent with the expanded proliferative and migratory keratinocyte compartment observed in Mlkl^-/-^ skin.

Further supporting this idea, treatment with necroptotic stimuli triggers the expression of differentiation markers in cultured keratinocytes. We found that primary mouse keratinocytes treated with a standard cell death assay dose of TSI had increased KRT1 expression as early as 2 hours post-treatment, comparable to the same cells exposed to a differentiation-inducing 0.65 mM calcium stimulus for 3 days (Fig. 6F). We also observed a time-dependent increase in expression of Keratin 1 and filaggrin in WT N/TERT cells treated with a moderate dose of TSI that was not observed in MLKL-deficient keratinocytes (Fig. 6G). Confocal imaging of WT N/TERT cells treated with TSI confirmed this increase at the single cell level, and an increase in the proportion of Keratin 1 positive cells that was not apparent in MLKL^-/-^ cells (Fig. 6H).

### N/TERT cells have increased calcium flux upon treatment with necroptotic stimuli

Using N/TERT cells that inducibly express GCaMP8 to monitor cytoplasmic calcium flux, we observed a dose-dependent increase in cytoplasmic calcium upon TSI treatment. Of note, while green fluorescence intensity due to Ca^2+^ flux was increased and sustained in cells in standard media (containing 0.15mM Ca^2+^), the effect could still be seen in cells treated in calcium-free media, with the medium to high dose TSI producing a similar degree of flux in the first 4 hours, regardless of extracellular calcium availability (Fig. 6I). This suggests that the initial calcium flux originates from intracellular stores, with extracellular replenishment contributing to maintaining the effect.

### CHMP4B overexpression reduces TSI-induced cell death

While N/TERT cells are resistant to TSI-induced cell death, we observed a small increase in dead cells at later time points with the highest TSI concentration. We also found that the absence of external calcium in the media increased N/TERT susceptibility to TSI-induced death (Fig. S6A), though notably never to a level comparable to HT29. We speculated that calcium-mediated membrane repair mechanisms may account for this difference, and based on work showing that the ESCRT protein CHMP4B co-localises with MLKL at the membrane during necroptosis and that extracellular Ca²⁺ was required for that translocation^29^, we tested the effect of overexpression of CHMP4B in WT N/TERT keratinocytes challenged with high-dose TSI ± external Ca^2+^.

We found that CHMP4B overexpression reduced minimal cell death in TSI-treated WT N/TERT cells to a similar extent under both standard and Ca^2+^-free conditions. However, in Ca^2+^-free conditions, the reduction in cell death was more modest relative to a higher baseline level of injury (Fig. 6J), consistent with our observation that N/TERT cells were more prone to TSI-induced death in Ca^2+^-free medium.

## Discussion

This study identifies a previously unrecognised function of necroptotic signalling in the skin. Rather than executing inflammatory cell death, the necroptotic machinery can act as a stress-adaptive program that diverts keratinocytes into a differentiated, non-lytic fate. Across diverse models of cutaneous injury, genetic or pharmacological inhibition of necroptosis accelerated wound closure. This indicates that RIPK3 and MLKL normally constrain the pace of repair, but via a cell death-independent mechanism.

MLKL has been reported to perform functions outside canonical necroptotic signalling, including roles in endosomal trafficking^30^, demyelination required for axon regeneration^31^, and regulation of steroidogenesis^32^, suggesting that it can operate independently of the canonical death pathway. Non-lytic outcomes arising from necroptotic signalling include enhanced extracellular vesicle release^30^, ESCRT-III–mediated membrane repair^29^ and calcium signalling^28,33^. These suggest that activation of RIPK3 and MLKL does not inevitably culminate in complete membrane rupture; however, the biological rationale for these sub-lytic states remains unclear. Our findings provide a mechanistic and physiological context for such activity in the skin, demonstrating how sub-lethal MLKL activation can drive keratinocyte differentiation.

Keratinocytes in intact skin are exposed to low levels of microbial-derived ligands^2,34^. Our data indicate that these signals primarily tune cellular sensitivity to death-inducing stimuli rather than directly drive tissue injury. Following barrier disruption, this input is amplified by increased exposure to microbial products, cytokines, and damage-associated signals, creating a trigger-rich environment that converges on the necroptotic pathway^1,5,12,13^. A lytic necroptotic response seems counterproductive in this setting, where it would amplify tissue damage without contributing to barrier restoration. Yet, the expression of RIPK3 and MLKL in basal keratinocytes and regenerating dermis indicates that the machinery for necroptotic signalling is present in intact skin, and even upregulated following damage and during tissue repair.

Furthermore, accelerated healing in the absence of RIPK3 or MLKL indicates that these proteins are functionally active. Still, the absence of necrotic keratinocytes at sites of re-epithelialisation in wild-type tissue indicates that this activity is non-lethal. Sub-lytic necroptotic signalling has been reported in other tissues, where it can lead to sustained inflammation^35^. In contrast, our data indicate that, in keratinocytes, necroptotic signalling is coupled to calcium flux-mediated differentiation and barrier repair, thereby providing a route to resolve the inflammatory sublytic state rather than sustain it.

Calcium is a central regulator of keratinocyte differentiation, with a well-described gradient across the epidermis and intracellular Ca²⁺ release acting as a key trigger for homeostatic differentiation^24,36^. Our data suggest that necroptotic signalling provides an alternative, stress-responsive route into this pathway. Activation of MLKL with TSI induced a dose-dependent rise in cytoplasmic Ca²⁺, with early flux occurring independently of extracellular calcium, consistent with release from intracellular stores. These findings indicate that sub-lethal MLKL activation can mobilise internal calcium to initiate differentiation, even in the absence of canonical extracellular calcium cues. Importantly, keratinocyte differentiation is accompanied by loss of necroptotic competence, with RIPK3 expression restricted to basal, proliferative keratinocytes. Diversion into differentiation may therefore not only prevent cell lysis but also extinguish further responsiveness to necroptotic stimuli.

While MLKL is most commonly associated with plasma membrane permeabilisation and lytic cell death, localisation to intracellular membranes has also been reported^37–39^. These observations have been difficult to reconcile with the prevailing view of MLKL as a terminal plasma membrane effector. Our data provide a context in which intracellular membrane engagement may be functionally meaningful. At higher levels of stimulation, Ca²⁺ dynamics reflected increasing membrane perturbation, with evidence of both intracellular release and leakage across the plasma membrane. Despite this, most cells remained viable, indicating that membrane repair mechanisms are likely engaged. Consistent with this, overexpression of the ESCRT-III component CHMP4B reduced cell death at higher TSI doses. Together, these observations indicate that MLKL activity engages both intracellular and plasma membrane compartments in keratinocytes. The internal calcium release can provide a route away from lysis by activating membrane repair programmes^40^ and potentially other calcium-mediated processes. Thus, MLKL-driven membrane permeabilisation may not be intrinsically lethal, at least in keratinocytes. Instead, our findings support a graded rheostat model in which the outcome of necroptotic signalling is shaped by the strength and duration of pathway engagement, the kinetics and extent of membrane perturbation, and the magnitude of calcium flux. How a cell interprets and resolves this signal is likely to depend on its intrinsic repair capacity, available fate programs, and tissue context. This framework may have broader relevance for understanding non-lytic outcomes of necroptotic signalling, but keratinocytes appear particularly well equipped to exploit it because calcium flux is already embedded in their terminal differentiation program.

It is well established that the normal Ca²⁺ gradient in the skin is lost upon barrier disruption^25^. MLKL-mediated mobilisation of intracellular calcium could therefore substitute for the extracellular calcium signals that are disrupted in wounded and disorganised skin. This may be especially relevant because restoration of the normal calcium gradient depends on barrier recovery^41^. In this context, necroptotic signalling may help restore aspects of barrier function by bypassing disrupted extracellular calcium cues, albeit at the cost of slower re-epithelialisation. Notably, MLKL deficiency does not impair normal skin development or steady-state barrier function^42^, and MLKL-deficient skin resolves to a fully functional barrier following injury. This suggests that the trade-off imposed by necroptotic signalling is specific to pathological contexts of barrier disruption, where MLKL-dependent differentiation may transiently favour structural reinforcement over rapid re-epithelialisation.

The adaptive value of this system is likely to be context-dependent and, in some settings, may be outweighed by competing demands. Notably, several mammalian lineages have independently lost key necroptotic components, including RIPK3 and MLKL^43–45^, indicating that this pathway is not universally required. These losses have largely been attributed to selective pressures imposed by pathogens, particularly viruses and intracellular bacteria, which encode factors that inhibit or subvert necroptotic signalling. In this view, pathogen-mediated inhibition of necroptosis may reduce its defensive value by allowing infected cells to remain viable and support pathogen replication^46–49^. Under these conditions, competing demands during tissue repair may exert greater selective influence. While our data cannot directly define the evolutionary pressures, they add additional context for interpreting these observations. Two examples highlight how distinct environmental pressures could shift this balance. Carnivores lack MLKL entirely^43^ and are commonly recognised to heal more rapidly than other mammals^50^. In animals that depend on predation, injury may compromise hunting and survival; thus, selective pressure may have favoured rapid wound closure in these species, at the expense of optimal barrier reconstruction. Cetaceans also appear to have lost MLKL function^43^. In these species, the surrounding aquatic environment reduces the requirement for a permeability barrier to prevent water loss^51,52^, diminishing the selective advantage of high-fidelity barrier restoration. In both cases, different pressures may converge on the same outcome: prioritising rapid wound closure over MLKL-dependent barrier reinforcement.

Translationally, our findings suggest that short-term, local modulation of necroptotic signalling could accelerate cutaneous repair by sustaining keratinocyte proliferation without risking long-term barrier disruption or systemic immune compromise. Direct modulation of RIPK3-MLKL activity may offer advantages over targeting individual upstream mediators, because necroptotic signalling integrates inflammatory cytokines, microbial signals, and tissue damage. This approach may be particularly valuable where rapid re-epithelialisation is clinically urgent, including in toxic epidermal necrolysis, and severe burns where epidermal barrier breakdown contributes to fluid loss, infection, and systemic complications^53,54^. Similarly, such inhibition may improve epidermal outgrowth from grafts, potentially increasing graft take and accelerating surface recovery in patients with severe burns.

Several limitations of our study remain. First, the upstream signals that engage RIPK3-MLKL signalling in vivo are not fully defined, including how caspase-8 activity is restrained within wounded skin. Second, our study does not directly resolve the molecular mechanisms linking MLKL activation to intracellular calcium mobilisation. Wound-associated metabolic rewiring may provide one possible explanation for the first of these gaps. The glycolytic protein PDK1 has been reported to promote RSK-dependent phosphorylation of caspase-8, reducing its destabilising effect on the necrosome^19^. It therefore represents a plausible mechanism by which hypoxic, glycolytic wound tissue^6,55^ could lower the threshold for RIPK3-MLKL activation in vivo. Although testing this mechanism directly is beyond the scope of this study, our observation that PDK1 is increased across models supports metabolic context as a future area of investigation. Similarly, while our data support a model in which sub-lethal MLKL activity perturbs intracellular membranes to release stored Ca²⁺, further work will be required to define the specific compartments involved and the mechanisms regulating this process. Dissecting how microbial priming, inflammatory gradients, metabolic cues, and MLKL-dependent calcium mobilisation are integrated will be important for understanding how necroptotic signalling is tuned between adaptive and lytic outcomes.

An additional limitation is the difficulty of directly establishing canonical necroptotic pathway activation in vivo. Expression of RIPK3 and MLKL does not, by itself, demonstrate pathway activation, and phosphorylated components remain the most direct markers of necroptotic engagement. However, reliable detection of endogenous pMLKL and pRIPK3 in tissue remains challenging, a limitation recognised across the field, with antibody specificity, fixation sensitivity, and context-dependent signal intensity limiting interpretation^56–58^. For this reason, we did not use pMLKL or pRIPK3 immunodetection to infer pathway activation in vivo. Instead, we took a rigorous genetic approach to assess pathway engagement, including the use of RIPK3 kinase-dead mice in addition to Ripk3^-/-^ and Mlkl^-/-^ mice. Similar phenotypes across genetic models support a role for canonical RIPK3-MLKL signalling. Furthermore, the cellular outcomes associated with defined necroptotic activation in vitro, including reduced proliferation and induction of keratinocyte differentiation, are recapitulated in vivo at sites of epidermal repair. While we cannot formally exclude the possibility that an alternative MLKL-dependent mechanism contributes to these effects in tissue, the convergence of genetic evidence and matched phenotypic outputs across systems strongly supports engagement of the canonical RIPK3-MLKL axis in vivo.

Finally, although our in vitro data demonstrate that human keratinocytes can undergo calcium-mediated differentiation in response to necroptotic stimulation, it remains to be determined whether comparable sub-lethal activation occurs in human skin in vivo. Establishing the presence and functional relevance of this adaptive version of the pathway in human wound healing will be an important step toward translating these findings into clinical applications.

## Methods

### Animal ethics and mouse strains

All mouse experiments were approved by the WEHI Animal Ethics Committee and conducted in accordance with the Australian Code for the Care and Use of Animals for Scientific Purposes. Mouse strains were generated and housed at the Walter and Eliza Hall Institute of Medical Research (WEHI) Specified Pathogen Free (SPF) Bioservices facilities in Parkville (VIC) 3052. Mice were maintained under constant temperature and humidity conditions on a 12 h light/dark cycle with ad libitum access to food and water. All strains were maintained on a C57BL/6 background and were backcrossed for a minimum of five generations.

Germ-free experiments were performed using C57BL/6 mice born and maintained in positive-pressure germ-free isolators. Experiments were conducted within the same isolators at WEHI’s germ-free facility in Kew (VIC) 3101.

Caspase-8^-/-^Ripk3^-/-^, Ripk1^+/-^Mlkl^-/-^^59^, Mlkl^-/-^Cflar^fl/f^(TgK14-CreERT2), Ifnγ^-/-^Tnf^-/-^, and FasL^gld/gld^Tnf^-/-^ compound mutant mice were generated at WEHI. Other mouse strains used in this study include Tnfr1^-/-60^, Ripk1^+/-61^, FasL^gld/gld62^, Tnf^-/-63^, Ripk1^D138N/D138N^ (kinase-dead)^22^, FasΔS^64^, FasΔM^64^, Ifnγ^-/-65^, Ripk3^-/-66^, Ripk3^D143N/D143N^ (kinase dead)^17^, Mlkl^-/-67^ Caspase-1^-/-68^, ASC^-/-69^, GsdmD^-/-70^, GsdmE^-/-71^, and Cflar^fl/f^Tg(K14-CreERT2)^20^.

### In vivo models

Smac-mimetic induced Toxic Epidermal Necrolysis (TEN): A TEN-like epidermal injury was induced by subcutaneous injection of 100μL of 1mg/mL of the Smac-mimetic (SM), Compound A (TetraLogic Pharmaceuticals, Malvern, PA, USA) formulated in 12% Captisol (CyDex Pharmaceuticals, Lawrence, KS, USA). In the standard screening protocol, male mice were injected in the right flank on day 0 and the left flank on day 2, providing two independent lesions per mouse. Mice were euthanised on day 3, yielding day 1 and day 3 lesions, and injection sites were photographed, clinically scored, and collected for ex vivo analyses.

Disease severity was assessed using a four-point ordinal scale (0–4) based on three macroscopic features associated with TEN: oedema, rubor, and epidermal disruption (Nikolsky sign/lesion severity). Each parameter was scored as 0 (not present), 1 (mild), 2 (moderate), or 3 (severe), and scores were summed to generate an overall clinical score^14,15^.

For selected strains, extended time-course experiments were performed with monitoring up to day 7 to assess lesion resolution. Based on accelerated healing observed in necroptosis-deficient strains in pilot experiments, additional day 5 endpoint cohorts were performed for these genotypes, with lesions scored as above and tissues collected for ex vivo analyses. In separate experiments, early time-course analyses were performed, with injection sites collected at 1, 3, or 6 h following SM administration to assess acute signalling events.

Statistical comparisons were performed using unpaired t-tests with Welch’s correction.

### Tamoxifen-induced epidermal-specific depletion of cFLIP

Keratinocyte-specific apoptotic injury was generated in Cflar^fl/fl^Tg(KRT14-CreERT2) mice by topical application of tamoxifen (2.5mg in 50µL 80% EtOH, 20% Lipoderm) to shaved dorsal skin, once daily for four consecutive days (day 0-3) to induce Cre-mediated recombination in basal keratinocytes. Cohorts were euthanised on days 4, 7, 9, and 11, sites were photographed, and tissue was collected for ex vivo analysis.

In the microbial-suppression study, 100mg of topical triple-antibiotic ointment (TAO, 400U Bacitracin Zinc, 3.5mg Neomycin Sulfate, 5000U Polymyxin B per gram cream in a Water/petroleum emulsion base) or carrier only was applied topically to the site, daily throughout the experiment. Microbial suppression was confirmed by plating skin swabs taken on day 7 onto agar and counting colony-forming units (CFU) following a 48-hour incubation at 37°C. Epidermal recovery was quantified at day 11 from images taken after scab removal and expressed as the percentage of the total application area with fully recovered epidermis.

### Full-thickness excision wound model

To induce mechanical injury, an area of dorsal skin was shaved and cleared with depilation cream under anaesthetic. Two 5 mm full-thickness wounds were generated on the cleared dorsal skin by placing the mouse on its side, pinching the dorsal skin at the midline and pinning it gently down, then punching through the two layers of skin with a sterile 5mm biopsy punch, leaving two full-thickness excision wounds on either side of the midline. Wounds were photographed for wound area assessment and covered with sterile transparent adhesive film dressings. Dressings were removed on day 3, to ensure that consistency in wound environments across cohorts was not affected by differences in dressing disruption by the mice. Wounds were photographed during recovery, and the wound area was calculated and expressed as a % relative to the original D0 wound. Cohorts were euthanised on days 6 or 8 post-wounding for ex vivo analysis.

For pharmacological inhibition of necroptosis, Nec-1s (30mg/kg) or vehicle control (25% PEG in PBS) was administered once daily IP throughout the experiment.

### Bone marrow transplantation

To determine the contribution of haematopoietic versus tissue compartments to the cutaneous-healing phenotypes, reciprocal bone marrow chimeras were generated between WT, Ripk3^−/−^, and Mlkl^−/−^ mice. Recipients were lethally irradiated (2 × 5.5 Gy, 4 h apart) and reconstituted the same day by intravenous injection of freshly isolated donor bone marrow (1 × 10^7 cells in 200 µL PBS). Mice were maintained on oral neomycin (1 g/L) for 3 weeks post-transplantation and housed for 16 weeks post-transplantation before use in TEN or excision wound assays. Chimerism was verified by WB on reconstituted bone marrow.

### Ex vivo Analyses

#### Histology and morphometric analysis

Skin samples were fixed in 10% neutral buffered formalin for 24-48 h then stored in 70% ethanol before paraffin embedding (FFPE) and sectioning. FFPE samples were sectioned for routine histology staining (haematoxylin and eosin; H&E) onto Superfrost slides (Thermo Fisher Scientific).

Epidermal thickness was measured as the average perpendicular distance between the epidermal base and the outer surface across at least 15 randomly selected sites per sample using ImageJ.

#### Immunohistochemistry

For immunohistochemistry (IHC) and immunofluorescence (IF), Superfrost Plus slides (Thermo Fisher Scientific, MA, USA) were used. Formalin-fixed, paraffin-embedded (FFPE) sections were dewaxed and rehydrated according to standard protocols. Slides were subjected to heat-induced antigen retrieval by boiling in citrate buffer for 20 min, rinsed in Tris-buffered saline containing 0.05% Tween-20 (TBST), then blocked and permeabilised in 3% normal goat serum (NGS) and 0.3% Triton X-100 in TBS for a minimum of 15 min.

For slides probed with anti-Filaggrin (BioLegend, San Diego, CA, USA) or anti-Keratin 1 (Biolegend), primary antibodies were diluted 1:500 in blocking buffer and applied to sections, which were incubated overnight at 4°C. The following day, slides were rinsed three times with TBST, blocked for endogenous peroxidase activity using Dako peroxidase block (Agilent Technologies, CA, USA) for 10 min, followed by avidin block and biotin block for 10 min each (Avidin/Biotin Blocking Kit; Vector Laboratories, MA, USA). Slides were then incubated with goat anti-rabbit biotinylated secondary antibody (Vector Laboratories) diluted 1:300 for 1 h, rinsed three times with TBST, incubated with VECTASTAIN Elite ABC HRP reagent (Vector Laboratories) for 30 min, rinsed, and developed using DAB substrate (Agilent Technologies). Slides were washed under running tap water, counterstained with haematoxylin, dehydrated through graded ethanol, cleared in xylene, and mounted using DPX mountant.

Slides stained for anti-cleaved caspase-3 (ASP175) (Cell Signalling Technology, MA, USA), anti-Ki67 (Thermo Fisher Scientific), anti-CD3 (Agilent Technologies), anti-CD45 (BD Biosciences), anti-F4/80 (WEHI in-house), H&E, and myeloperoxidase (MPO) were prepared using an Autostainer Link 48 (Agilent Technologies) according to standard protocols. Slides stained for anti-mRIPK3(8G7) (WEHI in-house; available from Merck Millipore as MABC1595), and anti-mMLKL(5A6) (WEHI in-house; available from Merck Millipore as MABC1634) were prepared according to optimised protocols^72^ using the Autostainer.

Entire slides were scanned with a Virtual Slide Microscope (Olympus) and viewed using CaseCenter (3DHISTECH, Budapest, Hungary) or scanned with a Polaris (PhenoImager HT) and viewed using QuPath v0.5.1 (open-source software, University of Edinburgh, Edinburgh, UK), and images were captured on a MacBook Pro 14-inch (Apple Inc., Cupertino, CA, USA). Minor image processing, such as adjustments to brightness and contrast, were applied to entire images, and the same adjustments were made to all comparable images.

### Whole-skin lysate preparation

Whole skin samples were lysed in death-induced signalling complex (DISC) lysis buffer by placing a piece of skin, approximately 100μg in 250μl of ice-cold DISC in a 2 mL Eppendorf tube (DISC lysis buffer comprised: 20 mM Tris pH 7.5, 2 mM EDTA, 1% Triton X-100, 150 mM sodium chloride, 10% glycerol, cOmplete cocktail protease inhibitor (Roche) in H2O). The samples were then finely diced in the tube with scissors and bead beat at 30 Hz for 3 x 1min using a TissueLyser (Qiagen; Venlo, Limburg, Netherlands). The samples were then lysed on ice for 30 min, spun at 13,000 rpm for 10 min, and the supernatant retained for protein analysis (ELISA, Bioplex and Western Blot). Protein lysates were quantified using a bicinchoninic acid (BCA) protein quantification kit (Thermo Fisher Scientific; MA, USA). Samples were normalised to the same concentration of total protein within an experiment.

### Cytokine analysis by ELISA and Bioplex

TNF, CCL2 and IL-6-ELISAs were performed using Invitrogen ELISA kits (Thermo Fisher Scientific) following the manufacturer’s protocols. Broader inflammatory profiling was performed using the Bio-Plex Pro™ Mouse Cytokine 23-Plex Assay (Bio-Rad). Results were normalised to total protein content and expressed as pg of target protein/mg total protein.

### Western blotting

Samples were normalised and equal amounts of protein were resolved by SDS–PAGE and transferred to PVDF membranes. Membranes were stained with Ponceau S to confirm uniform transfer and loading, and in most cases were imaged, and then blocked in 5% BSA or 5% skim milk. Ponceau S images were used as total-protein loading references for figure presentation. After destaining, membranes were incubated with primary antibodies overnight at 4 °C and detected with HRP-conjugated secondary antibodies using enhanced chemiluminescence (ECL). Where Ponceau S images were not available, membranes were subsequently probed with anti-β-actin and used as a loading control for figure presentation.

Primary antibodies used for Western blotting of mouse samples: anti-MLKL (clone 3H1; WEHI in-house; available from Merck Millipore as MABC604), anti-RIPK3 (clone 8G7; WEHI in-house; available from Merck Millipore as MABC1595), anti-HSP70 (clone N6; a gift from Dr R. Anderson, Peter MacCallum Cancer Research Institute, Melbourne, Australia), anti-cIAP1 (clone 1E1-1-10; Enzo Life Sciences), anti-PDK1 (clone D37A7; Cell Signalling Technology), anti-Keratin 1 (BioLegend), and anti-β-actin (Sigma-Aldrich).

### Cell culture and in vitro analysis

#### Keratinocyte culture

Human immortalised keratinocytes (N/TERT-1) and primary murine keratinocytes were cultured in KGM-Gold medium (Lonza, CC-3107) supplemented according to the manufacturer’s instructions and maintained at 37 °C with 5% CO₂. N/TERT cells were maintained in KGM-based medium without fetal calf serum and under low-calcium conditions (0.01 mM Ca²⁺). For calcium-depleted experiments, cells were washed twice and maintained in calcium-free KGM-Gold basal medium prior to stimulation. Cells were stimulated a minimum of 24 h after seeding. For live-cell imaging experiments, N/TERT cells were seeded in 96-well plates at 3,000 cells per well or in 24-well plates at 10,000 cells per well. For immunoblotting or immunofluorescence, cells were seeded into 12-or 6-well plates at approximately 30–40% confluency, depending on the assay.

### Induction of cell death signalling pathways

Necroptotic signalling was induced using TNFα, Smac-mimetic (SM; Birinapant or Compound A), and the pan-caspase inhibitor IDN-6556 (IDN), collectively referred to as TSI, at the following concentrations:

- Low (TSI-L): TNFα 4 ng/mL, SM 20 nM, IDN 2.5μM
- Medium (TSI-M): TNFα 25 ng/mL, SM 50 nM, IDN 2.5μM
- High (TSI-H): TNFα 100 ng/mL, SM 250 nM, IDN 5μM

ZBP1-dependent necroptotic signalling was induced by priming with IFNγ (20ng/mL) for 24h followed by treatment with Curaxin0137 (CBL; 5 μM) and IDN (2.5 μM).

### CRISPR/Cas9 genome engineering

CRISPR/Cas9-mediated editing was used to generate MLKL-knockout N/TERT cells and doxycycline-inducible reporter lines. Single-guide RNAs were cloned into a lentiviral CRISPR/Cas9 vector, and lentiviral particles were produced in HEK293T cells and used to transduce N/TERT cells. Following selection, single-cell clones were expanded and validated by sequencing and confirmed by absence of MLKL protein by immunoblotting.

For inducible GCaMP8m and CHMP4B expression, coding sequences were cloned into a lentiviral doxycycline-inducible expression vector. Stable lines were generated by lentiviral transduction and puromycin selection (1 μg/mL). Reporter expression was induced with doxycycline for 24 h prior to experiments and verified by fluorescence imaging (GCaMP8m) or immunoblotting (CHMP4B).

Due to the growth characteristics of N/TERT cells, expression of transgenes following lentiviral transduction typically became evident 6–10 days after infection.

To minimise exposure of N/TERT cells to high serum or calcium during lentiviral transduction, viral particles were generated in HEK293T cells using KGM medium supplemented with 1% fetal calf serum.

### Incucyte live cell imaging

Cell death and morphology were monitored using an IncuCyte S3 or S5X Live-Cell Analysis System (Sartorius). Plates were imaged using a 10× objective at 1 h intervals for 24–48 h. Cytotoxicity was quantified using propidium iodide (1:1000), SYTOX Green (1:10,000), or Cell Death NIR (1:2000), as indicated. The ratio of dye-positive cells to total cell area was calculated and normalised to baseline (0 h) per well.

For calcium flux analysis, doxycycline-inducible GCaMP8m-expressing N/TERT cells were induced for 24 h prior to stimulation. Cells were imaged at 1 h intervals using an IncuCyte S5X. Fluorescence intensity was quantified and normalised to baseline per well.

Quantification was performed using the Incucyte software prior to image export. Brightness and contrast adjustments were applied only to exported images used for figure presentation, were made uniformly across the entire image, and were matched across comparable images. Scratch assays were performed using IncuCyte ImageLock 96-well plates coated with extracellular matrix. N/TERT cells were seeded at 5,000 cells per well and grown to confluence. Wounds were generated using the IncuCyte 96-well WoundMaker tool (Sartorius). Detached cells were removed by washing twice with culture medium, and fresh medium ± TSI was added. Plates were placed immediately into an IncuCyte S3 Live-Cell Analysis System and imaged every 1–2 h for 24 h. Wound closure was quantified using the IncuCyte Scratch Wound Analysis Software Module and expressed as percentage closure relative to the initial wound area.

For all Incucyte experiments, quantification was performed using the Incucyte software prior to image export. Brightness and contrast adjustments were applied only to exported images used for figure presentation, were made uniformly across the entire image, and were matched across comparable images.

### Western blot analysis of cultured keratinocytes

For protein analysis, cells were lysed in RIPA buffer with protease and phosphatase inhibitors. Lysates were clarified, quantified by BCA assay, and processed for Western blotting as described for tissue samples. Membranes were stained with Ponceau S and imaged for use as loading controls.

Primary antibodies used for western blot of human keratinocytes: anti-MLKL (clone 3H1; WEHI in-house; available from Merck Millipore as MABC604), anti-phospho-MLKL (Ser358; Abcam; human-specific), anti-human RIPK3 (clone 1H2; WEHI in-house; human-specific available from Merck Millipore as MABC1640), anti-phospho-RIPK3 (Cell Signalling Technology; human-specific), anti-CHMP4B (Molecular Probes), anti-Keratin 1 (BioLegend, human specific), anti-filaggrin (BioLegend), and anti-ZBP1 (Thermo Fisher Scientific).

### Immunofluorescence

Cells were grown on collagen-coated 9mm round coverslips. Following experimental treatment cells were fixed in ice-cold 100% methanol, then blocked and permeabilised with 3% NGS and 0.3% Triton X-100 in TBS. Primary antibodies were applied overnight at 4 °C, followed by Alexa Fluor-conjugated secondaries. Nuclei were counterstained with DAPI.

Primary antibodies used for immunofluorescence: anti-Keratin 1 (BioLegend, human-specific). Immunofluorescence images were acquired on a Zeiss LSM 980 confocal microscope and processed using ImageJ/Fiji (NIH, Bethesda, MD, USA). Image processing included maximum-intensity Z-projection and minor linear adjustments to brightness and contrast. Adjustments were applied to entire images, with channel-specific settings held constant across comparable samples. Images were recoloured to colourblind-friendly palettes using Adobe Photoshop (Adobe Inc., San Jose, CA, USA).

### Declaration of generative AI and AI-assisted technologies in the writing process

During the preparation of this work, the authors used ChatGPT (OpenAI) and Claude AI (Anthropic) to improve the readability, clarity, and structure of the text, reduce the word count, and identify editorial errors. After using these tools, the authors reviewed and edited the content as needed and take full responsibility for the content of the published article.

### Author contributions

Conceptualisation, H.A. and J.S.; Methodology, H.A., Y.H., A.L.S., and W.C.; Investigation, H.A., Y.H., N.S., A.L.-G., G.L.H., S.B., K.S., E.B.-S., A.H., and W.C.; Formal Analysis, H.A. and Y.H.; Resources, J.M.M and A.L.S.; Writing – Original Draft, H.A.; Writing – Review and Editing, H.A., J.S., J.M.M, A.L.S, E.B-S; Visualisation, H.A. and Y.H.; Supervision, H.A., J.S., A.L.S., and J.M.M; Funding Acquisition, H.A., J.S and J.M.M.

## Acknowledgements

Funding sources: This work was supported by National Health and Medical Research Council (NHMRC) Investigator Grants awarded to H.A. (grant number 1194144), J.M.M. (grant number 2034104) and J.S. (grant numbers 1195038 and 2034661). H.A. and J.S. received philanthropic funding support from Thomas William Francis & Violet Coles Trust, facilitated by George Varigos. Y.H. is supported by a Leo Foundation Scholarship administered through Melbourne Health. S.B. is supported by the CareerTrackers Indigenous Internship Program at WEHI. J.M.M. and J.S. have received research funding from Anaxis Pharma Pty Ltd, and A.H.’s salary was paid by Anaxis Pharma Pty Ltd. This work was made possible through the Victorian State Government Operational Infrastructure Support Program and the Australian Government NHMRC IRIISS.

We are grateful to the WEHI laboratories and researchers who generously provided mouse strains used in this study. In particular, we thank Seth Masters for Asc^-/-^ and Caspase-1^-/-^ mice, Lorraine O’Reilly for FasΔM and FasΔS mice, James Vince for Gsdmd^-/-^ and Gsdme^-/-^ mice, and Chris Tonkin for Ifng^-/-^mice.

The authors gratefully acknowledge the WEHI Centre for Dynamic Imaging, the WEHI Advanced Histotechnology Facility and WEHI Bioservices for their support and assistance in this work.

We also thank Robin Anderson, Peter MacCallum Cancer Research Institute, Melbourne, Australia for the HSP-70 Mab.

## Declaration of Interests

Y.H. receives scholarship support from the Leo Foundation. A.H., A.L.S., J.M.M. and J.S. contribute to, or have contributed to, a program to develop necroptosis pathway inhibitors in collaboration with Anaxis Pharma Pty Ltd. All other authors declare no interests.

**Supplementary Figure S1.**

(A) Control data for Fig 1C - Epidermal thickness of untreated skin from mice of the indicated genotypes. N=4-7, each data point is the average of ≥ 10 measurements per animal. Box plots show the interquartile range (25th–75th percentile) with the centre line indicating the median and whiskers representing minimum and maximum values. (B) Immunohistochemistry of WT skin following SM injection showing Ki67, cleaved caspase-3 (CC3), CD3, and CD45 at untreated baseline (UT) and 1, 3, and 6 hours post-injection. Scale = 100μM. (C) CCL2 and TNF concentrations in skin lysates from the indicated genotypes in UT, day 1, and day 3 lesional sites following SM injection, measured by ELISA. N=4-10, each data point is a separate animal; bars are mean ± SEM. (D) Quantification of wound width at day 7 in WT, Mlkl^-/-^, and Ripk3^-/-^ mice. Measured on H&E as the distance from healing edge to healing edge. N=6-9, each data point is a separate animal; bars are mean ± SEM. (E) Heatmap showing row-normalised Z-scores of multiplex cytokine measurements for the indicated mediators in whole skin lysates from WT, Mlkl^-/-^, and Ripk3^-/-^ mice at UT and day 1 following SM injection. Each row is scaled independently to highlight relative changes across conditions. Alongside are absolute cytokine concentrations (pg/mg tissue) for selected analytes from the multiplex panel. N=3, each data point is a separate animal; bars are mean ± SEM.

Supplementary Figure S2.

(A) Example of a mild responder among germ-free (GF) mice following SM injection. Representative clinical images at days 1 and 3 and corresponding cleaved caspase-3 (CC3) IHC at day 1 are shown. Scale =100μM.

Supplementary Figure S3

(A) Immunohistochemistry for CD3, Myeloperoxidase (MPO), and F4/80 in skin from cFlip^K14ERcre^ and Mlkl^-/-^cFlip^K14ERcre^ mice at untreated baseline (UT) and days 7, 9, and 11 counted from the start of the 4-day tamoxifen treatment.

(B) Heatmap showing row-normalised Z-scores of multiplex cytokine measurements in whole skin lysates from cFlip^K14ERcre^ and Mlkl^-/-^cFlip^K14ERcre^mice from days 4, 7, 9, and 11 following the start of tamoxifen treatment. Each row is scaled independently to highlight relative changes across conditions. Absolute cytokine concentrations (pg/mg tissue) for selected analytes from the multiplex panel on the right-hand side. N=6-8, each point represents an individual animal, bars are mean ± SEM.

(C) Immunoblot analysis of PDK1 in whole skin lysates from cFlip^K14ERcre^ and Mlkl^-/-^cFlip^K14ERcre^ following tamoxifen-induced epidermal cFLIP deletion. Ponceau staining is shown as a loading control.

**Supplementary Figure S4**

(A) Immunohistochemistry for RIPK3 and MLKL in wound edges from WT and Mlkl^-/-^ mice.

(B) Immunoblot analysis of whole-skin lysates from WT mice during wound healing (day 0, day 1, day 3, day 6, and day 8) showing MLKL expression. Ponceau staining is shown as a loading control.

(C) Immunoblot analysis of whole-skin lysates from WT, Mlkl^-/-^, and Ripk3^-/-^ mice at day 0, day 3, and day 6 following wounding. Ponceau staining is shown as a loading control.

**Supplementary Figure S5.**

(A) Single agent controls for Figure 5D. Relative cell growth measured as confluence over time in N/TERT cells treated with TNF, SMAC mimetic, or IDN alone. N = 3 independent experiments (n=3 wells per experiment).

(B) Quantification of cell death in N/TERT and HT29 cells following induction of ZBP1-dependent necroptosis using interferon priming, CBL0137, and caspase inhibition (ICI), also showing IFNγ and CBL single agent controls. Measured as dead cells per image, normalised to 0h timepoint (n = 3 technical replicates).

(C) Representative phase-contrast images of WT and Mlkl^-/-^ N/TERT cells untreated (UT) or treated with interferon, CBL0137, and caspase inhibition at 0, 4, 12, and 24 hours. Arrows indicate example cells with differentiating morphology. Wide image scale = 400μM, enlarged image scale = 200μM. Arrows indicate example cells with differentiated morphology.

Data on graphs are shown as mean ± SEM.

**Supplementary Figure S6 legend**

(A) Quantification of cell death in WT N/TERT keratinocytes treated with increasing concentrations of TSI in standard (0.15mM Ca^2+^) or calcium-free medium. Representative experiment shown with n=3 technical replicates, plotted mean ± SEM.

