## Supplementary figures and images for "Necroptotic Signalling Diverts Keratinocyte Fate to Promote Differentiation and Slow Wound Healing"

### Supplementary Figures S1-S6

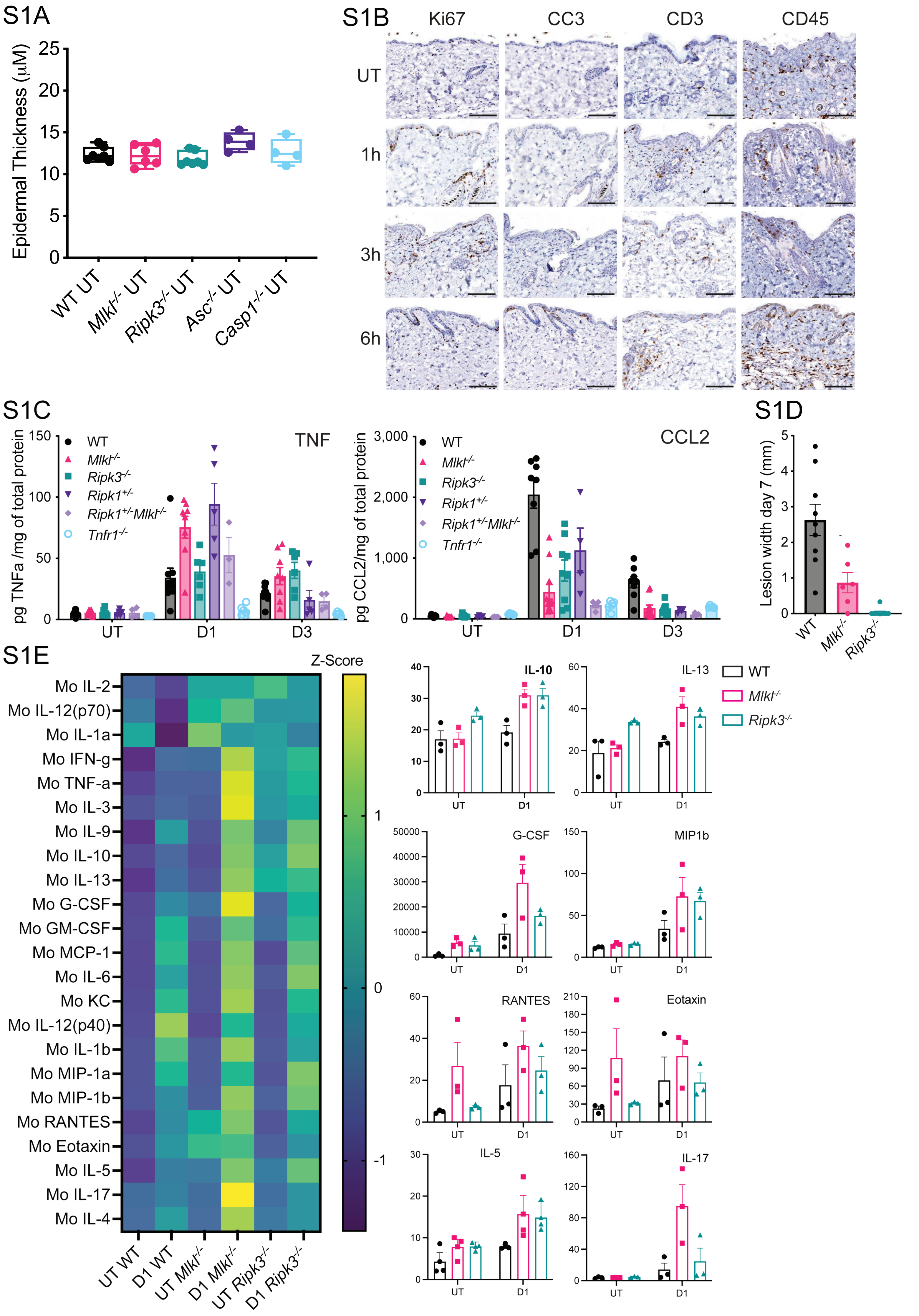

S2A

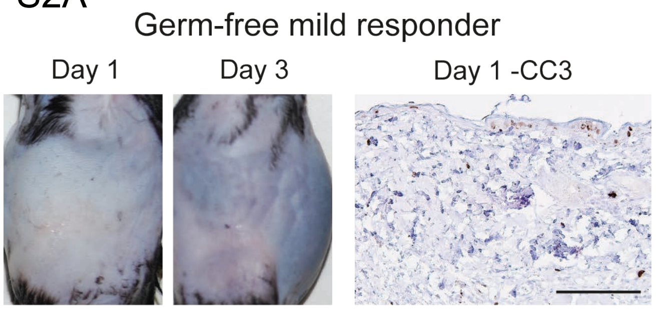

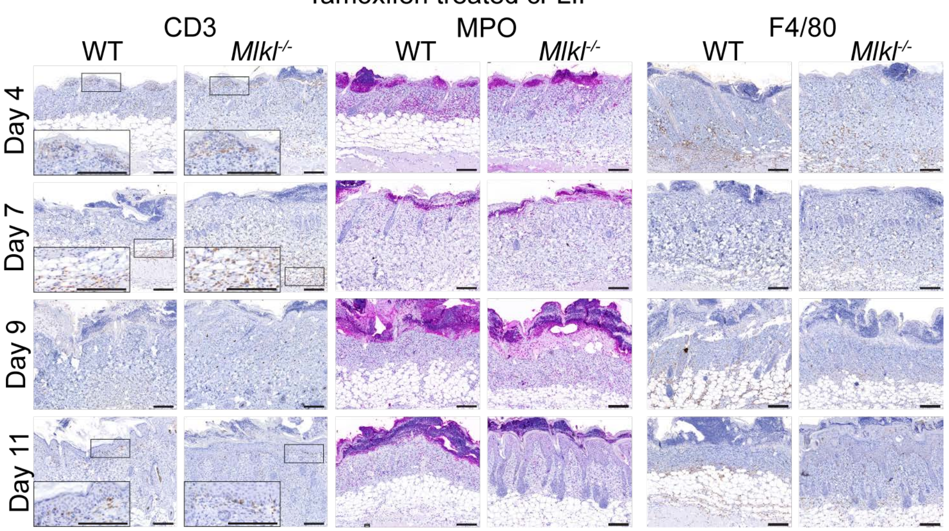

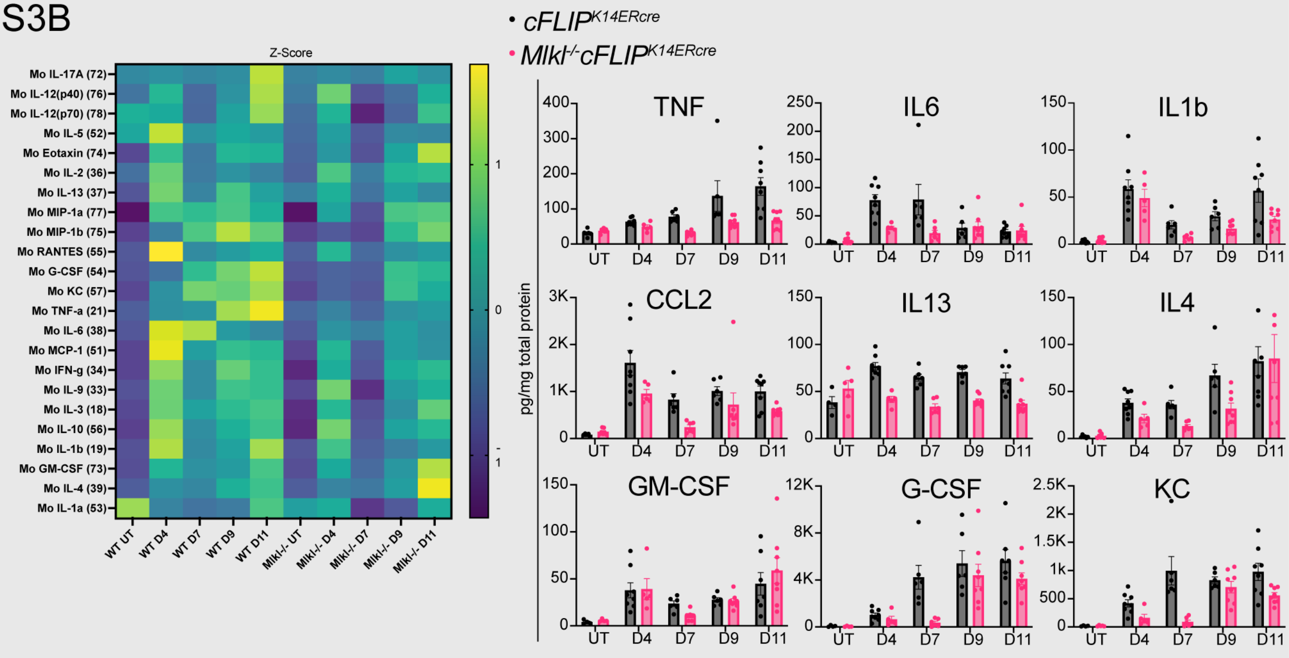

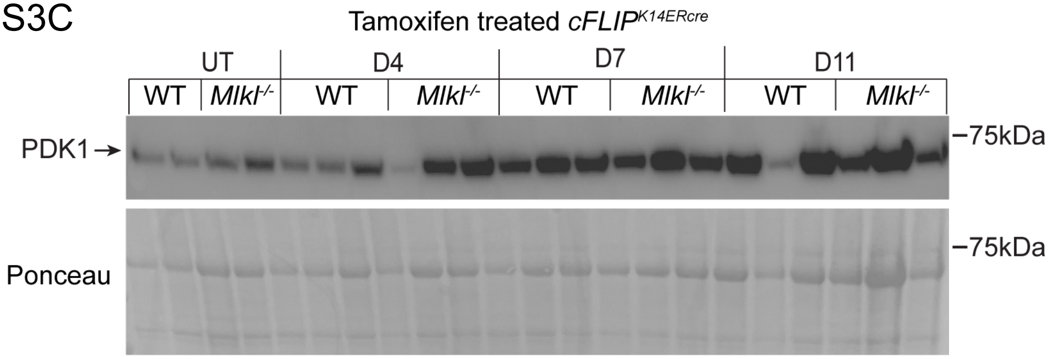

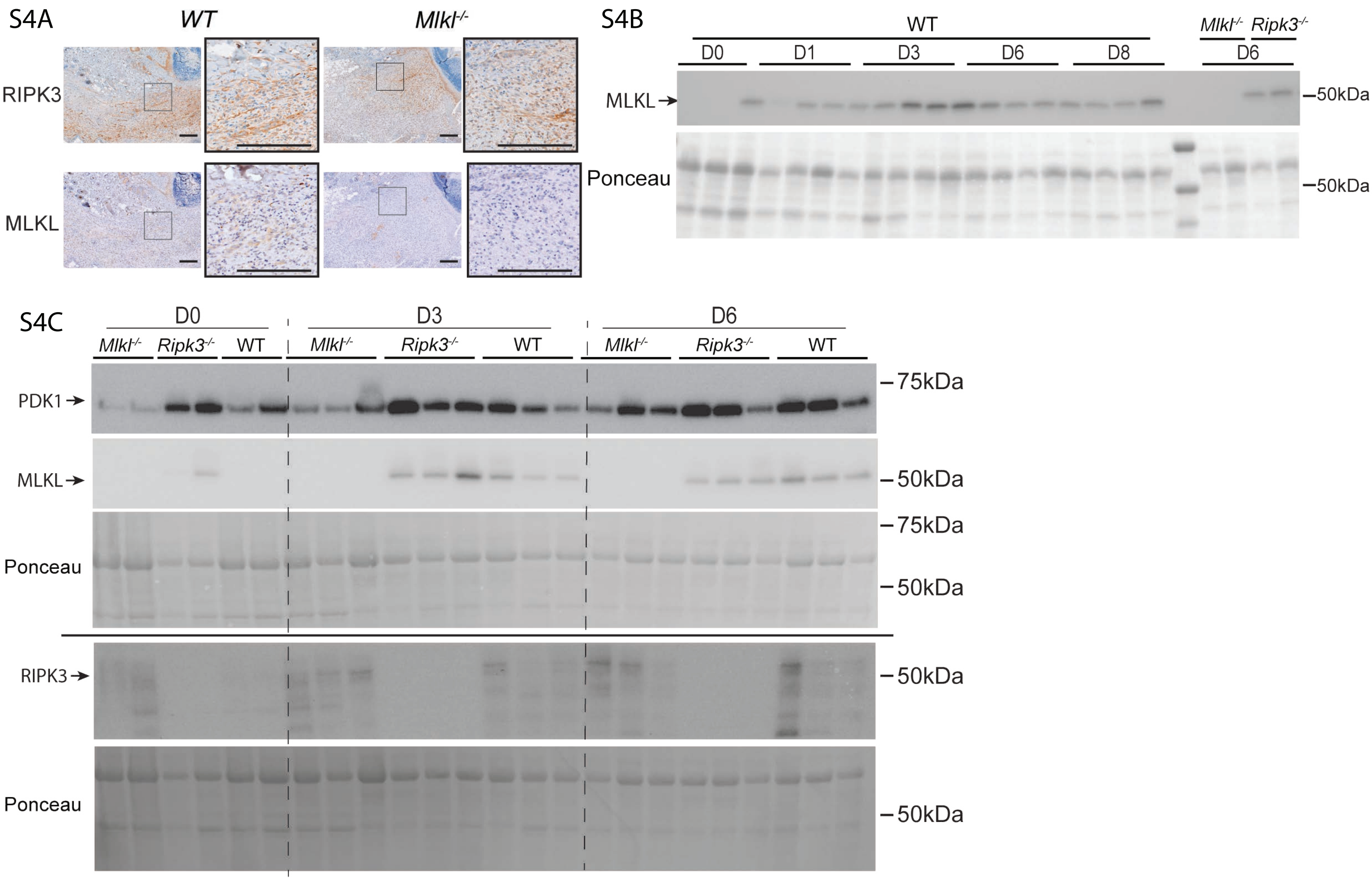

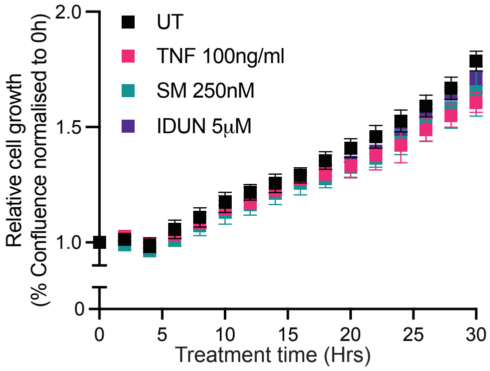

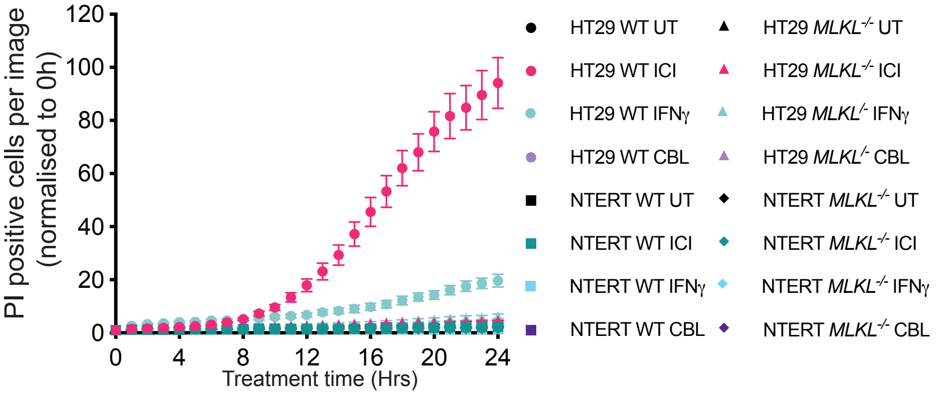

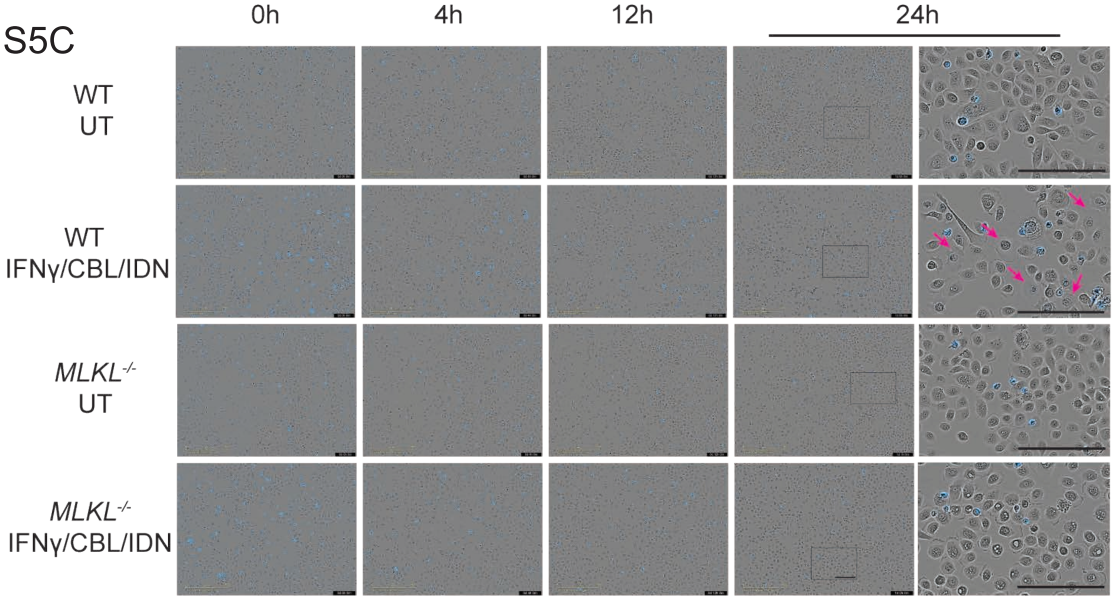

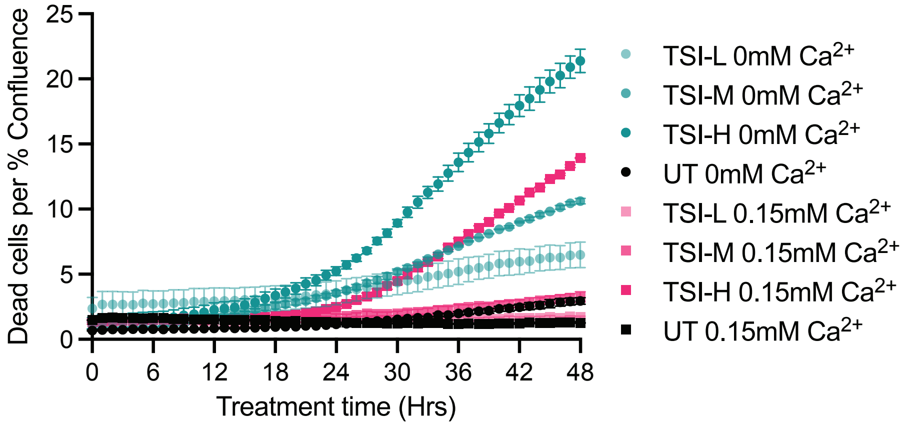
