## Supplementary Table S1 for "Necroptotic Signalling Diverts Keratinocyte Fate to Promote Differentiation and Slow Wound Healing"

| Row Statistics Figure 1A | **Day 1** | | | **Day 3** | | | **Day 5** | | |
| --- | --- | --- | --- | --- | --- | --- | --- | --- | --- |
|  | **Mean** | **SD** | **N** | **Mean** | **SD** | **N** | **Mean** | **SD** | **N** |
| ***WT*** | 6.51 | 1.01 | 35 | 7.27 | 1.46 | 33 | 6.23 | 1.48 | 13 |
| ***Tnfr1^-/-^*** | 0.40 | 0.55 | 5 | 0.60 | 0.55 | 5 |  | | |
| ***Ripk1^+/-^*** | 1.40 | 0.55 | 5 | 1.29 | 0.49 | 7 |  |  |  |
| ***Ifnγ^-/-^Tnf^-/-^*** | 1.70 | 0.82 | 10 | 1.45 | 0.93 | 11 |  |  |  |
| ***FasL^gld/gld^Tnf^-/-^*** | 1.20 | 0.45 | 5 | 1.40 | 0.55 | 5 |  |  |  |
| ***Casp8^-/-^Ripk3^-/-^*** | 1.00 | 0.71 | 5 | 2.20 | 0.84 | 5 |  |  |  |
| ***FasL^gld/gld^*** | 1.40 | 0.55 | 5 | 2.40 | 1.14 | 5 |  |  |  |
| ***Tnf^-/-^*** | 2.18 | 0.98 | 11 | 1.93 | 1.00 | 14 |  |  |  |
| ***Ripk1^KD/KD^*** | 2.00 | 0.82 | 4 | 2.25 | 0.96 | 4 |  |  |  |
| ***Ripk1^+/-^Mlkl^-/-^*** | 3.00 | 0.82 | 4 | 1.50 | 1.00 | 4 |  |  |  |
| ***Fas^ΔS^*** | 3.57 | 1.27 | 7 | 3.43 | 1.27 | 7 |  |  |  |
| ***Ifnγ^-/-^*** | 4.20 | 1.03 | 10 | 3.70 | 2.00 | 10 |  |  |  |
| ***Ripk3^KD/KD^*** | 6.11 | 1.17 | 9 | 4.17 | 0.75 | 6 | 1.75 | 0.50 | 4 |
| ***Ripk3^-/-^*** | 6.00 | 1.18 | 14 | 2.25 | 1.16 | 8 | 1.00 | 1.26 | 6 |
| ***Mlkl^-/-^*** | 6.25 | 1.36 | 12 | 7.50 | 0.84 | 6 | 3.50 | 1.52 | 6 |
| ***Fas^ΔM^*** | 6.25 | 0.75 | 12 | 7.17 | 0.83 | 12 |  | | |
| ***Casp1^-/-^*** | 5.80 | 1.10 | 5 | 7.20 | 0.45 | 5 |  |  |  |
| ***Asc^-/-^*** | 6.20 | 1.48 | 5 | 6.60 | 1.14 | 5 |  |  |  |
| ***Gsdmd^-/-^*** | 6.40 | 0.89 | 5 | 6.80 | 1.30 | 5 |  |  |  |
| ***Gsdme^-/-^*** | 6.60 | 1.82 | 5 | 7.40 | 0.55 | 5 |  |  |  |
